# HNF1A is a Novel Oncogene and Central Regulator of Pancreatic Cancer Stem Cells

**DOI:** 10.1101/238097

**Authors:** Ethan V. Abel, Masashi Goto, Brian Magnuson, Saji Abraham, Nikita Ramanathan, Emily Hotaling, Anthony A. Alaniz, Chandan Kumar-Sinha, Michele L. Dziubinski, Sumithra Urs, Lidong Wang, Jiaqi Shi, Meghna Waghray, Mats Ljungman, Howard C. Crawford, Diane M. Simeone

## Abstract

The biological properties of pancreatic cancer stem cells (PCSCs) remain incompletely defined and the central regulators are unknown. By bioinformatic analysis of a PCSC-enriched gene signature, we identified the transcription factor HNF1A as a putative central regulator of PCSC function. Levels of HNF1A and its target genes were found to be elevated in PCSCs and tumorspheres, and depletion of HNF1A resulted in growth inhibition, apoptosis, impaired tumorsphere formation, PCSC depletion, and downregulation of OCT4 expression. Conversely, HNF1A overexpression increased PCSC numbers and tumorsphere formation in pancreatic cancer cells and drove PDA cell growth. Importantly, depletion of HNF1A in primary tumor xenografts impaired tumor growth and depleted PCSCs *in vivo*. Finally, we established an HNF1A-dependent gene signature in PDA cells that significantly correlated with reduced survivability in patients. These findings identify HNF1A as a central transcriptional regulator of the PCSC state and novel oncogene in pancreatic ductal adenocarcinoma.

## INTRODUCTION

Pancreatic ductal adenocarcinoma (PDA) is projected to be the 2^nd^ leading cause of cancer deaths in the U.S. by 2020 (Rahib et al., 2014). The exceeding lethality of PDA is attributed to a complex of qualities frequent to the disease including early and aggressive metastasis and limited responsiveness to current standards of care. While both aspects are in-and-of-themselves multifaceted and can be partially attributed to factors such as the tumor microenvironment (Olive et al., 2009, Provenzano et al., 2012, Waghray et al., 2016) and the mutational profile of the tumor cells (Yachida et al., 2012), cancer stem cells (CSCs) have also been identified to contribute to promoting early metastasis and resistance to therapeutics (Hermann et al., 2007, Li et al., 2011).

CSCs, which were originally identified in leukemias (Bonnet and Dick, 1997, Graham et al., 2002), have been identified in a number of solid tumors including glioblastoma (Singh et al., 2003), pancreas (Li et al., 2007, Hermann et al., 2007) and colon (O’Brien et al., 2007). In these cases, CSCs have been characterized by the ability to establish disease in immunocompromised mice, to resist chemotherapeutics, the capability of both self-renewal and differentiation into the full complement of heterogeneous neoplastic cells that comprise the tumor, and the propensity to metastasize. In each case, CSCs are distinguished from other tumor cell types by the expression of various, sometimes divergent cell surface markers. Our lab was the first to identify pancreatic cancer stem cells (PCSCs), which were found to express the markers EPCAM (ESA), CD44, and CD24 (Li et al., 2007). In addition to these markers, CD133 (Hermann et al., 2007), CXCR4 (Hermann et al., 2007), c-MET (Li et al., 2011), aldehyde dehydrogenase 1 (ALDH1) (Kim et al., 2011), and autofluorescence (Miranda-Lorenzo et al., 2014) have all been proposed markers of PCSCs. In all cases, the identified cells are characterized by being able to form spheres of cells (tumorspheres) under non-adherent, serum-free conditions, as well as an increased ability to form tumors in mice compared to bulk tumor cells. While a number of markers have been identified for PCSCs, relatively little is known about the transcriptional platforms that govern their function and set them apart from the majority of bulk PDA cells. Transcriptional regulators such NOTCH (Wang et al., 2009, Abel et al., 2014), BMI1 (Proctor et al., 2013), and SOX2 (Herreros-Villanueva et al., 2013) have been demonstrated to play roles in PCSCs, although these proteins are also critical for normal stem cell function in many tissues.

In this study we sought to better understand the biological heterogeneity of PCSCs and their bulk cell counterparts in an effort to identify novel regulators of PCSCs in the context of low-passage, primary patient-derived PDA cells. Using microarray analysis and comparing primary PDA cell subpopulations with different levels tumorigenic potential and stem cell-like function, we identified hepatocyte nuclear factor 1-alpha (HNF1A), an endoderm-restricted transcription factor, as a key regulator of the PCSC state. Supporting this hypothesis, depletion of HNF1A resulted in a loss of PCSC numbers and functionality both *in vitro* and *in vivo*. Additionally, ectopic expression of HNF1A augmented PCSC properties in PDA cells and enhanced growth and anchorage-independence in normal pancreatic cell lines. Mechanistically, we found that HNF1A regulates the stem cell transcription factor OCT4 (POU5F1), which is necessary for stemness in PCSCs. Based on these data we postulate a novel pro-oncogenic function for HNF1A through its maintenance of the pancreatic cancer stem cell state.

## RESULTS

### An HNF1A gene signature dominates a PCSC gene signature

A transcriptional profile of PCSCs has yet to be established, and we hypothesized that such a profile would contain key regulators of the PCSC state. To pursue this hypothesis, we utilized a series of low-passage, patient derived PDA cell lines to isolate PCSC-enriching and non-enriching subpopulations for comparative analysis. Using two of our previously described PCSC surface markers, CD44 and EPCAM (Li et al., 2007), we found that low-passage PDA cells generally formed three subpopulations (abbreviated P herein) based on surface staining: CD44^High^/EPCAM^Low^ (P1), CD44^High^/EPCAM^High^ (P2), or CD44^Low^/EPCAM^High^ (P3) (Supplementary Figure 1A). Similar expression patterns were observed in 10 primary tumor samples (data not shown). Using previously described measures of PCSC function (Li et al., 2007, Li et al., 2011), including co-expression of the PCSC marker CD24 (Supplementary Figure 1B), the ability for isolated subpopulations to reestablish cellular heterogeneity (Supplementary Figure 1C), the ability to form tumorspheres under non-adherent/serum-free culture conditions (Supplementary Figure 1D), and initiate tumors in immune-deficient mice (Supplementary Table 1), we found that P2 cells showed greater enrichment for cells with PCSC properties than their P1 and P3 counterparts.

Using 2 primary PDA lines (NY8 and NY15), P1, P2, and P3 PDA cells were sorted by flow cytometry, prepped immediately for mRNA, and analyzed by Affymetrix GeneChip microarray and validated by quantitative RT-PCR. We found that P2 cells from both lines exhibited a signature of 50 genes that was upregulated (>1.5 fold) relative to both P1 and P3 cell counterparts (Figure 1A). To further refine this gene cohort, we utilized oPOSSUM (Kwon et al., 2012), a web-based system to detect overrepresented transcription factor binding sites in gene sets. Interestingly, HNF1A, a P2 cohort gene itself (Figure 1B), had predicted binding sites in the ±5000 regions (from start of transcription) of 17/50 of the enriched genes, and due to its stringent consensus sequence (DGTTAATNATTAAC) was the most highly ranked common transcription factor by Z-score (17.895). Of these 50 genes, HNF1A is known to positively regulate cohort genes *HNF4A* (Boj et al., 2001), *NR5A2* (Molero et al., 2012), *CDH17* (Zhu et al., 2010), *IGFBP1* (Babajko et al., 1993, Powell and Suwanichkul, 1993), and *DPP4* (Gu et al., 2008). Interestingly, genome-wide association (GWA) studies have recently identified certain single nucleotide polymorphisms (SNPs) in the HNF1A locus as risk factors for developing PDA (Pierce and Ahsan, 2011b, Li et al., 2012, Wei et al., 2012), although the mechanism by which these SNPs exert their influence is currently unknown. To further support the enrichment of HNF1A in PCSCs, sorted cells were western blotted for HNF1A and one of HNF1A’s target proteins, CDH17. Both proteins were found to be elevated in P2 cell lysates compared to other subpopulations (Figure 1C), in agreement with their transcript levels. Additionally, surface expression of DPP4 (CD26) was also found to be highest on cells in the P2 subpopulation (Figure 1D). CSCs are enriched in cancer cell populations grown under low-attachment tumorsphere (S) conditions compared to cells grown in adherent (A) conditions. In keeping with this observation, we found protein levels of HNF1A and CDH17 elevated in multiple PDA lines cultured under tumorsphere conditions (Supplementary Figure 2A, data not shown). Using a GFP-based reporter driven by 8 tandem copies of the HNF1A consensus sequence GGTTAATGATTAACC (Supplementary Figure 2B), we found GFP expression was elevated in NY5, NY8, and NY15 cells grown under tumorsphere (S) compared to adherent conditions (A) (Supplementary Figure 2C). This construct showed excellent dependence on HNF1A expression as targeting HNF1A with an HNF1A-specific siRNA ablated expression of both the ectopic GFP and endogenous CDH17 (Supplementary Figure 2D). Lastly, we found the frequency of GFP positive cells increased in cells grown in suspension, with GFP expression being highest in the P2 subpopulation of NY15 cells (Supplementary Figure 2E). Based on our gene expression and tumorsphere data, we hypothesized that HNF1A is a central regulator of CSC function.

**Figure 1:**
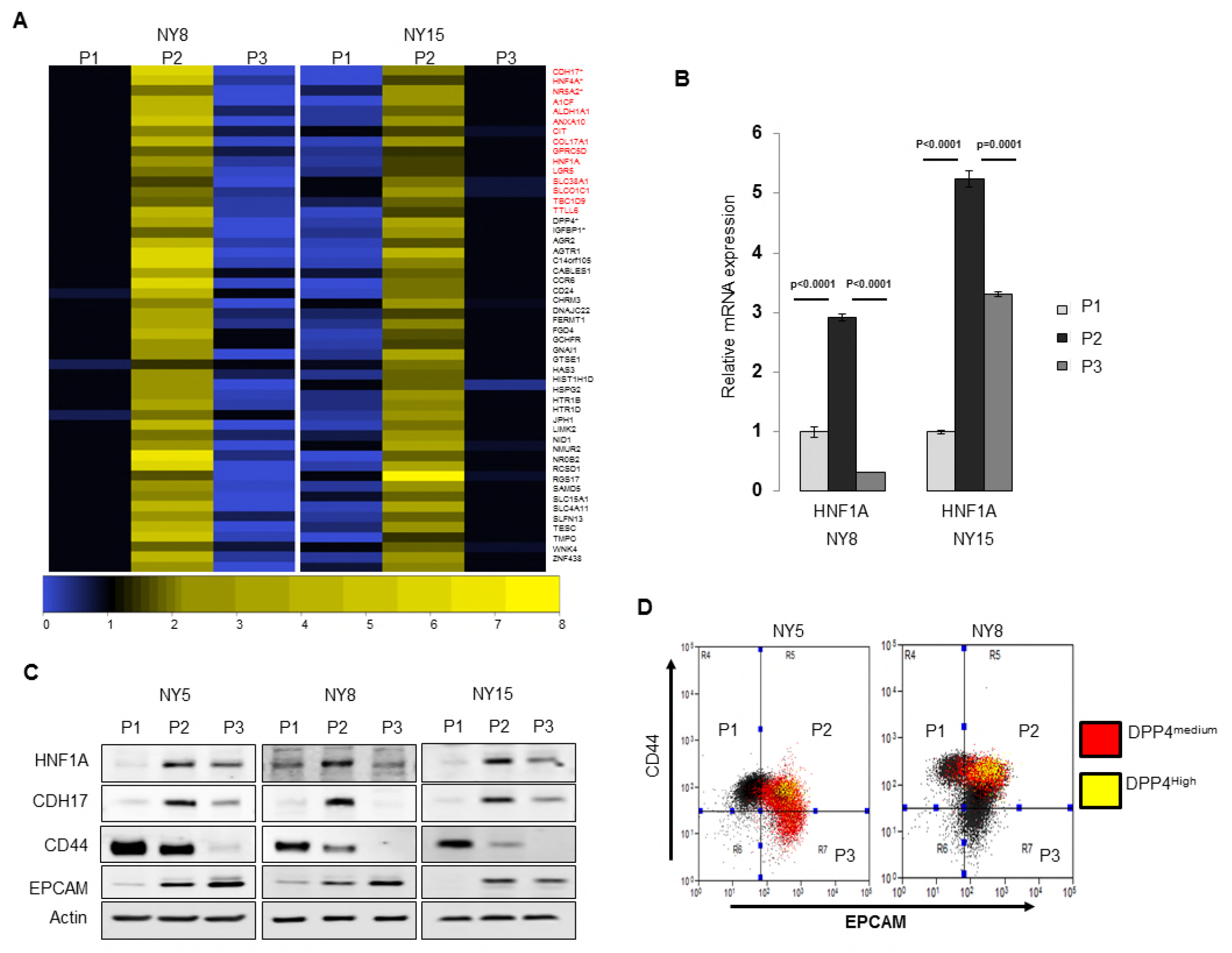
HNFlA-signature dominates pancreatic CSCs. (A) Heatmap representing relative fold differences in qRT-PCR expression of 50 cancer stem cell-enriched genes in NY8 and NY15 cells. Per-gene values are relative to P1 or P3, whichever is higher. Gene names in red text indicate predicted HNF1A targets and asterisks (*) indicate known HNF1A targets. P1: EPCAM^Low^/CD44^High^, P2: EPCAM^High^/CD44^High^, P3: EPCAM^High^/CD44^Low^. For all genes, expression levels were normalized to an *ACTB* mRNA control, n=3 biological replicates. Only genes with a significant (p<0.05) increase in P2 over both P1 and P3 subpopulations are shown. (B) qRT-PCR analysis of *HNF1A* mRNA expression, normalized to an *ACTB* mRNA control, from different primary PDA subpopulations (n=3 biological replicates). Statistical difference was determined by one-way ANOVA with Tukey’s multiple comparisons test; ***p<0.001, ****p<0.001. (C) Western blot analysis of HNF1A and target gene CDH17, as well as CD44 and EPCAM as subpopulation controls in sorted PDA cells. (D) Co-localization of HNF1A target gene DPP4 surface expression with EPCAM and CD44 expression is highlighted in red (moderate expression) and yellow (high expression). Related data can be found in **Supplementary Figures 1 and 2**.

**Figure 2:**
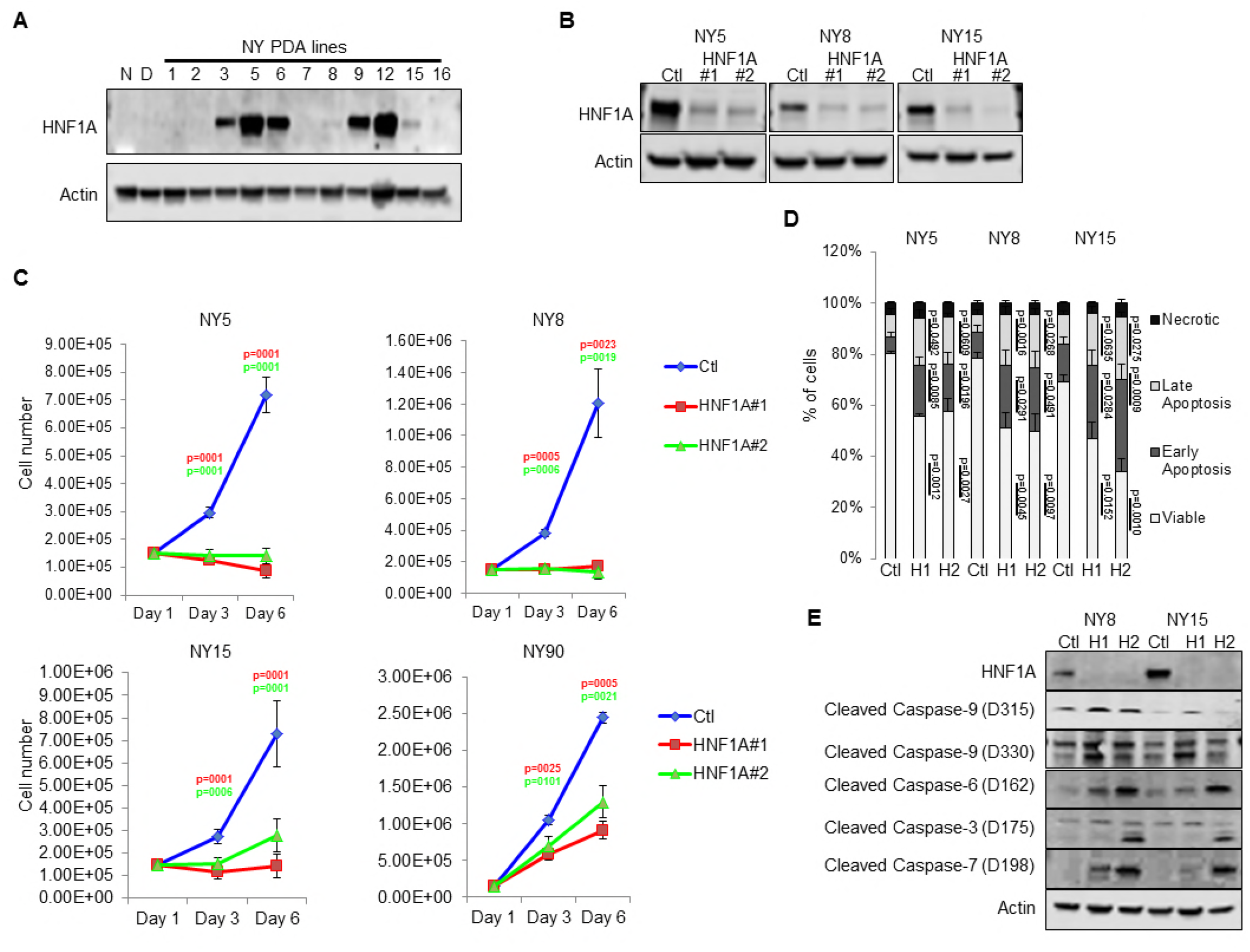
Knockdown of HNF1A in primary PDA cells inhibits growth in vitro. (A) Western blot analysis of HNF1A expression in a panel of primary PDA lines compared to immortalized pancreatic ductal cell line HPNE (N) and HPDE (D). (B) Western blot of NY5, NY8, NY15 cells transfected with non-targeting (Ctl) or HNF1A-targeting siRNA for 3 days, showing effective depletion of HNF1A protein by RNAi. (C) 1.5x10^5^ PDA cells were transfected with control (Ctl) or HNF1A-targeting siRNA. Cells were collected and manually counted 3 and 6 days after transfection (n=3 biological replicates). Statistical difference was determined by one-way ANOVA with Dunnett’s multiple comparisons test. Red and green p values indicate Ctl vs. HNF1A#1 or #2, respectively. (D) Annexin V/DAPI staining was performed on NY5, NY8, and NY15 cells transfected with control (Ctl) or HNF1A-targeting siRNA (H1, H2) for 3 days. The amount of viable (annexin V-/DAPI-), early apoptotic (annexin V+/DAPI-), late apoptotic (annexin V+/DAPI+), and necrotic (annexin V-/DAPI+) cells are quantitated (n=4 biological replicates). Statistical difference was determined by one-way ANOVA with Dunnett’s multiple comparisons test, with p values indicated to the right of each subpopulation relative to the control subpopulations. (E) Western blot analysis of cleaved caspases in NY8 and NY15 cells following HNF1A-knockdown (3 days). Actin serves as a loading control. Related data can be found in **Supplementary Figure 3**.

### HNF1A is a critical regulator of CSC function in PDA cells

Consistent with our hypothesis that HNF1A may be an integral component of PDA biology we observed higher levels of HNF1A protein and transcripts in PDA cells compared to nontransformed immortalized pancreatic cell lines HPNE (N) and HPDE (D) (Figure 2A; Supplementary Figure 3A, B). Immunostaining of a PDA tissue microarray showed HNF1A expression to be significantly elevated (p<0.0001) in PDA neoplastic ducts (n=41) compared to normal pancreatic ducts (n=18) (Supplementary Figure 3C, D). To examine the role of HNF1A in PDA cells, we depleted the protein with two distinct siRNAs (Figure 2B). Knockdown of HNF1A resulted in reduced cell numbers in several primary PDA lines (Figure 2C).

Interestingly, knockdown of HNF1A resulted in more profound growth inhibition in cells with moderate to high HNF1A expression (NY5, NY8, NY15) than cells with low HNF1A expression (NY90) (Figure 2A, C; Supplementary Figure 3A, B). To determine whether the apparent loss in cell number was due to apoptotic cell death, we performed annexin V/DAPI staining on control and HNF1A-depleted NY5, NY8, and NY15 cells. In all cases, knockdown of HNF1A resulted in a significant (p<0.05) increase in early and late apoptotic cells, while not affecting necrotic cell numbers (Figure 2D). Furthermore, increased cleavage of caspases 3, 6, 7, and 9 was observed in cells depleted of HNF1A (Figure 2E), indicating apoptotic cell death. These data indicate that HNF1A is important for PDA cell growth/survival and basal expression levels predict response to targeting.

Next we pursued whether depletion of HNF1A impacted PDA subpopulation distribution. Consistent with a central role in maintaining PDA cell heterogeneity, we consistently found that knockdown of HNF1A in multiple PDA lines resulted in loss of P2 cells (Figure 3A), supporting a role for HNF1A in maintaining PDA cell heterogeneity. In addition to a decrease in CD44 surface expression, we also observed a marked decrease in CD24 surface expression (Figure 3B, C) and mRNA levels (data not shown); suggesting that loss of HNF1A depletes the CSC compartment. Knockdown of HNF1A did not change surface expression levels of EPCAM (Figure 3A, data not shown). To assess functional consequences of HNF1A-depletion on the PCSC compartment, cells (NY8, NY15) expressing HNF1A shRNAs were grown under tumorsphere-promoting conditions. These shRNAs effectively depleted HNF1A as well as CDH17 (Figure 3D), indicating downstream signaling inhibition. Consistent with a role in PCSC function, HNF1A knockdown showed a marked reduction in tumorsphere formation (p<0.05) (Figure 3E, F).

**Figure 3:**
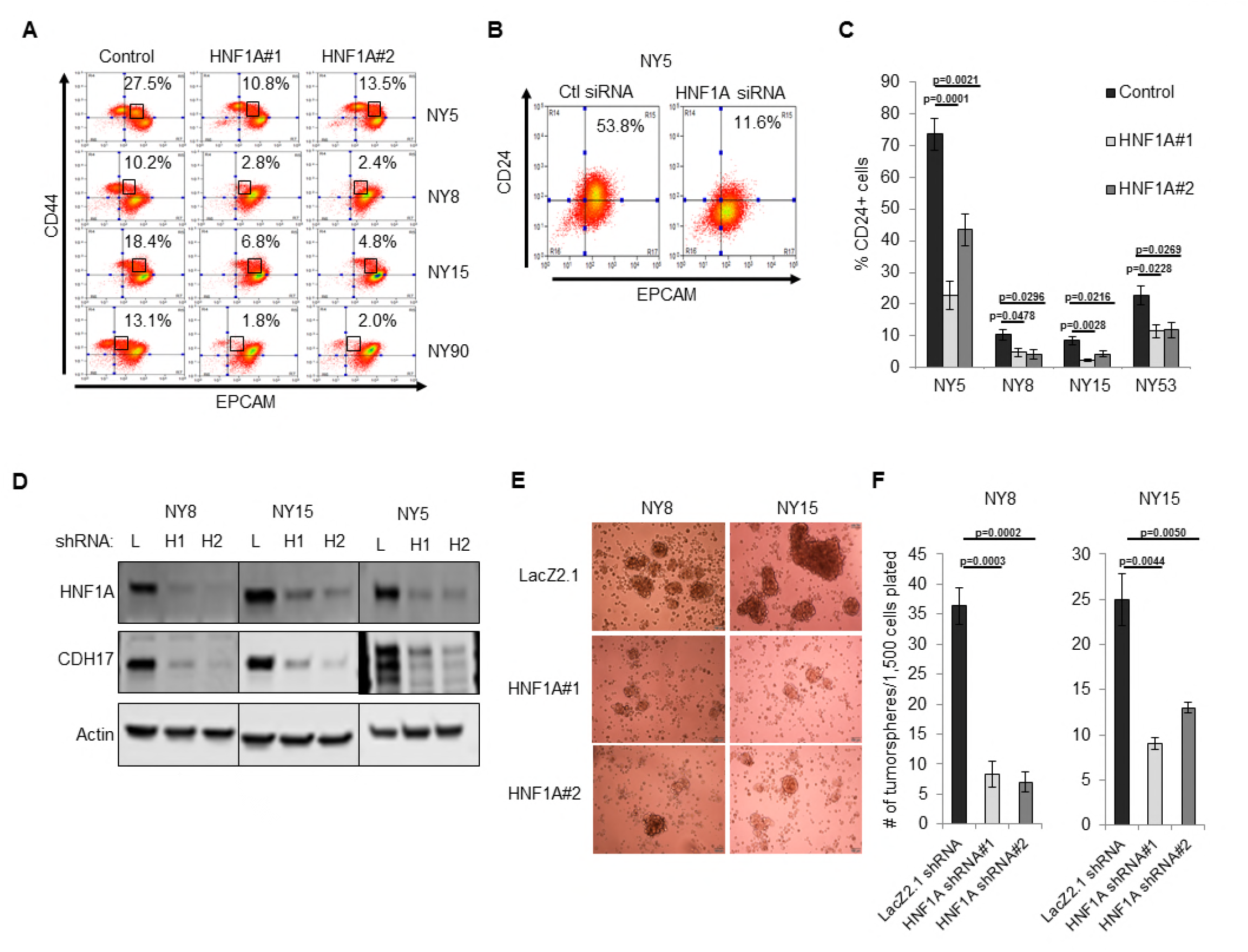
Knockdown of HNF1A depletes CSC numbers. (A) Surface expression of CD44 and EPCAM in multiple primary PDA cells following HNF1A knockdown. Percentage of P2 cells are indicated. (B) Representative CD24 and EPCAM surface staining in cells following HNF1A knockdown (6 days) (C) Quantitation of CD24+ cells in multiple primary PDA cells following HNF1A knockdown, n=4 biological replicates. Statistical difference was determined by one-way ANOVA with Dunnett’s multiple comparisons test. (D) NY8, NY15, and NY5 cells expressing LacZ2.1 (L) or two distinct HNF1A-targeting shRNAs (H1 and H2) were lysed and western blotted for HNF1A, CDH17, and Actin, showing effective knockdown of HNF1A and downstream signaling (CDH17). (E, F) NY8 and NY15 cells expressing LacZ2.1 or HNF1A-targeting shRNAs were grown in tumorsphere media on non-adherent plates (1500 cells/well). The number of tumorspheres formed after 6 days were counted (n=3 biological replicates). Representative images of spheres (100X magnification) are shown in (F) with quantitation in (E). Statistical difference was determined by one-way ANOVA with Dunnett’s multiple comparisons test.

### HNF1A exhibits oncogenic properties in pancreatic cells

We next sought to determine whether CSC properties could be augmented by ectopic expression of HNF1A in PDA cells. Using doxycycline-inducible expression of HNF1A (Figure 4A, B), we noted increased expression of CD24, CD44, and EPCAM in multiple primary PDA lines (Figure 4B-D, data not shown). To determine whether HNF1A could drive tumorsphere formation, using NY15 and NY53 (a model for low endogenous HNF1A expression) we found that HNF1A-expressing cells formed ~2.5 fold more tumorspheres than their counterparts (Figure 4E). Similar results were also seen in moderate-HNF1A expressing cells NY8 (data not shown).

We next examined the effects of ectopic HNF1A expression in the non-tumorigenic pancreatic ductal cell lines HPDE and HPNE, which were devoid of endogenous HNF1A expression (Figure 2A). Doxycycline-inducible ectopic expression of HNF1A alone or in concert with ectopic KRAS^G12D^ was readily achieved in HPDE cells (Supplementary Figure 4A). Consistent with previous reports, KRAS^G12D^ induced phosphorylation of both ERK1/2 and AKT in HPDE cells. Similar effects were seen in HPNE cells constitutively expressing HNF1A and KRAS^G12D^ alone or in combination (Supplementary Figure 4A). We then tested the impact the of HNF1A and/or KRAS^G12D^ expression, either alone or in combination, on HPDE cell growth. Under normal growth conditions with serum, (LacZ) HPDE cells grew to confluency but did not form colonies, presumably due to contact-inhibition (Supplementary Figure 4B). Expression of KRAS^G12D^, however, resulted in colony formation, indicating a bypass of contact inhibition. HNF1A alone resulted in significantly increased colony formation, which was further enhanced by the additional expression of KRAS^G12D^. Similar effects were seen in HPNE cells (data not shown). In clonogenicity assays, HNF1A-expressing HPNE cells formed similar numbers of colonies to control and KRAS^G12D^-expressing cells (Figure 4F, G), however, HNF1A alone promoted enhanced colony size. HPDE cells failed to form colonies at clonal densities in the presence of serum. In addition to foci formation, anchorage-independent growth can indicate cellular transformation *in vitro*. When suspended in soft agar, control HPDE cells failed to grow over a 21-day period (Figure 4H, I). The addition of KRAS^G12D^ alone did not significantly promote colony formation, consistent with its relatively weak transforming ability in HPDE cells. Interestingly, HNF1A alone resulted in numerous small colonies which in turn synergized with the expression of KRAS^G12D^ in the form of numerous large colonies. Neither HNF1A nor KRAS^G12D^ alone resulted in anchorage-independent growth in HPNE cells (data not shown). Lastly, we examined the effects of both transgenes on PCSC marker expression. Expression of HNF1A increased expression of EPCAM, CD44, and CD24 in HPDE cells (Supplementary Figure 4A, C). Control HPNE cells lacked expression of both EPCAM and CD24, but expressed high levels of CD44. Expression of HNF1A was able to increase CD44 surface expression, while not changing EPCAM status (Supplementary Figure 4C, data not shown). Remarkably, CD24 was potently induced upon HNF1A expression, with nearly 84% of HPNE cells expressing CD24 compared to 0.5% of LacZ-expressing control cells. These data would suggest that HNF1A possesses properties of an oncogene capable of cooperation with oncogenic KRAS. *HNF1A is required for tumor growth and cancer stem cells maintenance in vivo* To determine whether HNF1A was necessary for tumorigenesis, we implanted two primary lines (NY5 and NY15) expressing control or two HNF1A-targeting shRNAs orthotopically in the pancreas of NOD/SCID mice. HNF1A-depleted cells showed significantly reduced tumor growth compared to their control cohorts (p<0.05), (Figure 5A, B). Similar results were observed with HNF1A knockdown in subcutaneous xenografts of NY 5 and NY15 cells (Figure 5C, Supplementary Figure 5A). To determine whether inhibition of tumor growth was due to effects on the PCSC compartment, NY5 tumors were dissociated and analyzed by flow cytometry. Consistent with our *in vitro* findings, the EPCAM+/CD44+/CD24+ cell population was significantly reduced in HNF1A-depleted tumors (p<0.05) (Figure 5D, E). Importantly, western blot analysis of resultant tumor lysates confirmed that shRNAs remained effective at depleting HNF1A during the course of the experiment (Supplementary Figure 5B). Interestingly, Mason’s trichrome staining of NY5 tumors showed marked increase in mouse stromal infiltration and greater epithelial cell architecture in HNF1A-depleted tumors compared to control tumors (Supplementary Figure 5C), the latter suggesting a shift to a more differentiated tumor histology.

**Figure 4:**
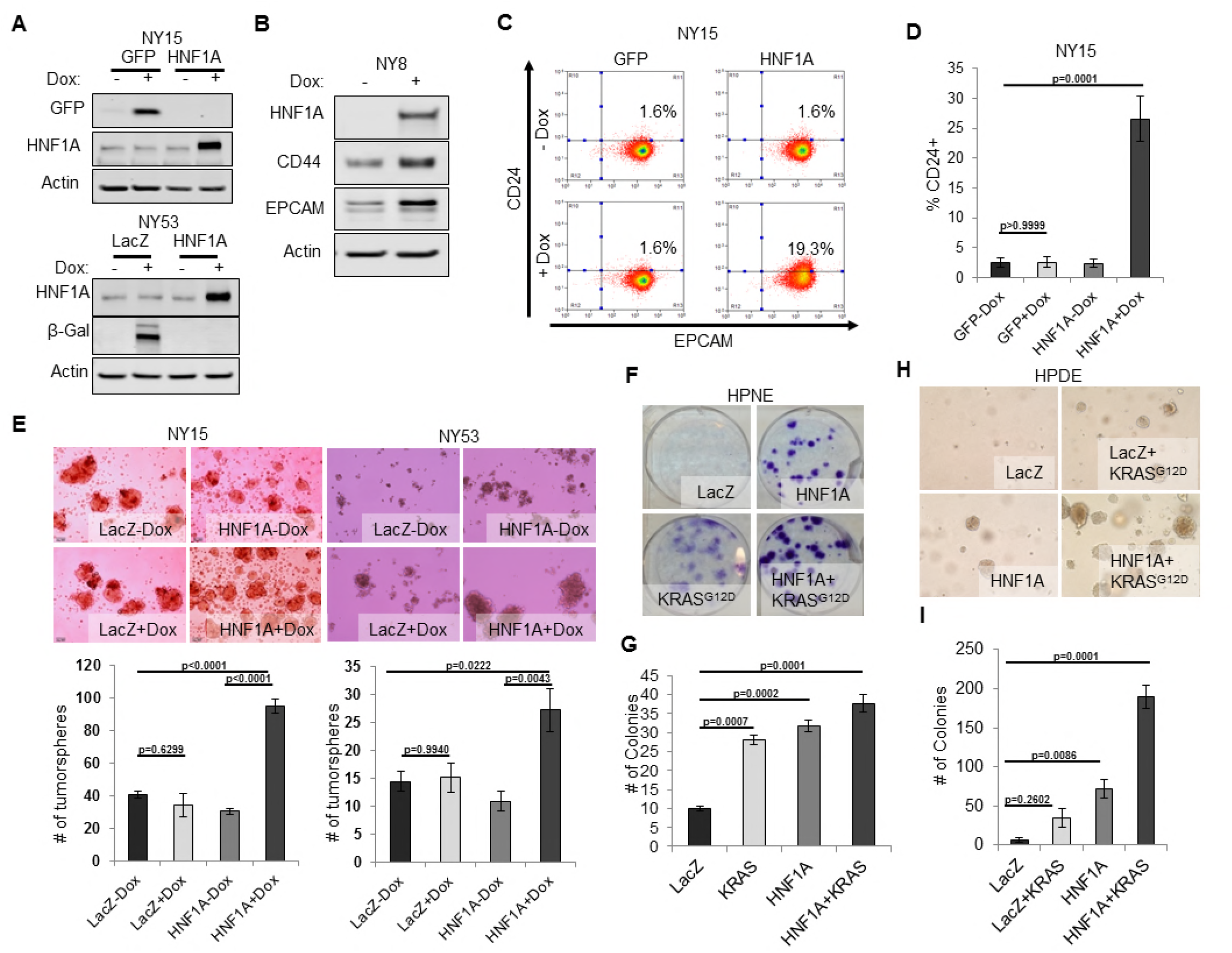
Overexpression of HNF1A promotes CSC properties in PDA cells and normal pancreatic cell lines. (A) NY15 and NY53 cells Western blotted for HNF1A and control gene induction 48 hrs ± doxycycline (Dox). (B) NY8 cells were treated 48 hrs ± Dox to induce ectopic HNF1A. Lysates were western blotted for HNF1A, Actin, and PCSC markers EPCAM and CD44. (C) Representative surface expression of CD24 and EPCAM on NY15 cells expressing GFP or HNF1A. (D) Quantitation of CD24+ NY15 GFP and HNF1A cells by flow cytometry (n=3 biological replicates). Statistical difference was determined by one-way ANOVA with Tukey’s multiple comparisons test. (E) NY15 and NY53 LacZ and HNF1A cells were grown under sphere-forming conditions ± Dox. The number of tumorspheres formed after 7 days were quantitated (n=4 biological replicates). Statistical difference was determined by one-way ANOVA with Tukey’s multiple comparisons test. Representative images (100X magnification) of spheres are shown in the lower panels. (F, G) HPNE LacZ and HNF1A cells were plated at 200 cells/6cm dish and treated ± Dox for 2 weeks, fixed, and stained with crystal violet (F). (G) Resultant colonies were quantitated (n=3 biological replicates). Statistical difference was determined by one-way ANOVA with Dunnett’s multiple comparisons test. (H, I) HPDE cells expressing inducible LacZ, LacZ with KRAS^G12D^, HNF1A, or HNF1A with KRAS^G12D^ were embedded in soft agar + Dox and monitored for signs of anchorage-independent growth for 21 days. (H) Representative images of resultant colonies (100X magnification) and (I) quantitation of colonies after 21 days (n=3 biological replicates). Statistical difference was determined by one-way ANOVA with Dunnett’s multiple comparisons test. Related data can be found in **Supplementary Figure 4**.

### HNF1A regulates stemness through OCT4 expression

As a direct relationship between HNF1A and stem cell function has not been reported, we examined mRNA expression of central stemness regulators *MYC, SOX2, KLF4, NANOG*, and *POU5F1* (OCT4) in HNF1A-depleted cells. Of these transcription factors, only *POU5F1* mRNA showed consistent downregulation in multiple PDA cell lines in response to HNF1A knockdown (Figure 6A, data not shown). Similarly, we found that *POU5F1* mRNA was upregulated in response to overexpression of HNF1A in both PDA cells and HPDE cells (Figure 6B), indicating regulation of OCT4 expression by HNF1A in pancreatic-lineage cells. OCT4 has previously been shown to be elevated in PCSCs (Miranda-Lorenzo et al., 2014, Luo et al., 2017), although a functional role for the protein has not been demonstrated in this context. To determine if OCT4 regulation was a key event in HNF1A-dependent stemness, we targeted OCT4 with multiple siRNA, either in combination or as single sequences. Depletion of OCT4 resulted in a pronounced inhibition of tumorsphere formation, comparable to HNF1A knockdown (Figure 6C-G). Importantly, knockdown of either HNF1A or OCT4 had comparable effects on the protein levels of OCT4A (Figure 6C), the isoform responsible for imparting stemness (Lee et al., 2006). To determine whether expression of OCT4A was sufficient to rescue stemness of PDA cells depleted of HNF1A, NY8 and NY15 cells were transduced with OCT4A-expressing lentiviruses or vector controls and transfected with HNF1A siRNA. Consistent with our previous results, loss of HNF1A impaired tumorsphere formation in both lines expressing the vector control, however, this effect was overcome by the expression of OCT4A (Figure 6H, Supplementary Figure 6A, B). These data indicate that HNF1A mediates stemness of PCSCs through regulation of OCT4. *An HNF1A gene signature is associated with poor survival in PDA patients* Lastly we sought to gain insight into the transcriptional activity of HNF1A in PDA and determine whether its transcriptome held prognostic information similar to other signatures in PDA (Bailey et al., 2016, Collisson et al., 2011). In order to identify transcriptional targets of HNF1A, we performed Bru-seq, a variation of RNA-seq which measures changes in nascent RNA levels *(bone fide* transcription rate) as opposed to steady-state RNA changes measured by conventional RNA-seq and microarray (Paulsen et al., 2013). Using concomitant ChIP-seq from control and HNF1A depleted NY8 and NY15 cells using an HNF1A-specific antibody, we identified 243 HNF1A-activated and 46 HNF1A-repressed transcripts shared between NY8 and NY15 (Figure 7A). 139/239 (57.2%) and 11/46 (23.9%) HNF1A-activated/repressed genes showed detectable HNF1A binding (Figure 7B), either distal or proximal to the transcriptional start site, supporting the role of HNF1A as a transcriptional activator. Importantly, a number of known HNF1A target genes exhibited HNF1A promoter-proximal binding and transcriptional responsiveness via Bru-seq/ChIP-seq, including *CDH17* (Figure 7C). Additionally, the PCSC marker *EPCAM* also showed HNF1A distal binding and transcriptional responsiveness, implicating HNF1A as a direct regulator of this gene. *CD24*, which showed changes in transcription in response to HNF1A loss, did not show direct binding, indicating an indirect mechanism of regulation (data not shown). *POU5F1* transcription was found to decrease in both NY8 (34.3%) and NY15 (41.5%) cells, with weak enrichment of a putative regulatory region previously shown to bind HNF1A (Consortium, 2012, Malakootian et al., 2017) (data not shown). These data suggest that while *POU5F1* transcription is promoted by HNF1A, the mode of regulation may be a combination of direct and indirect mechanisms.

**Figure 5:**
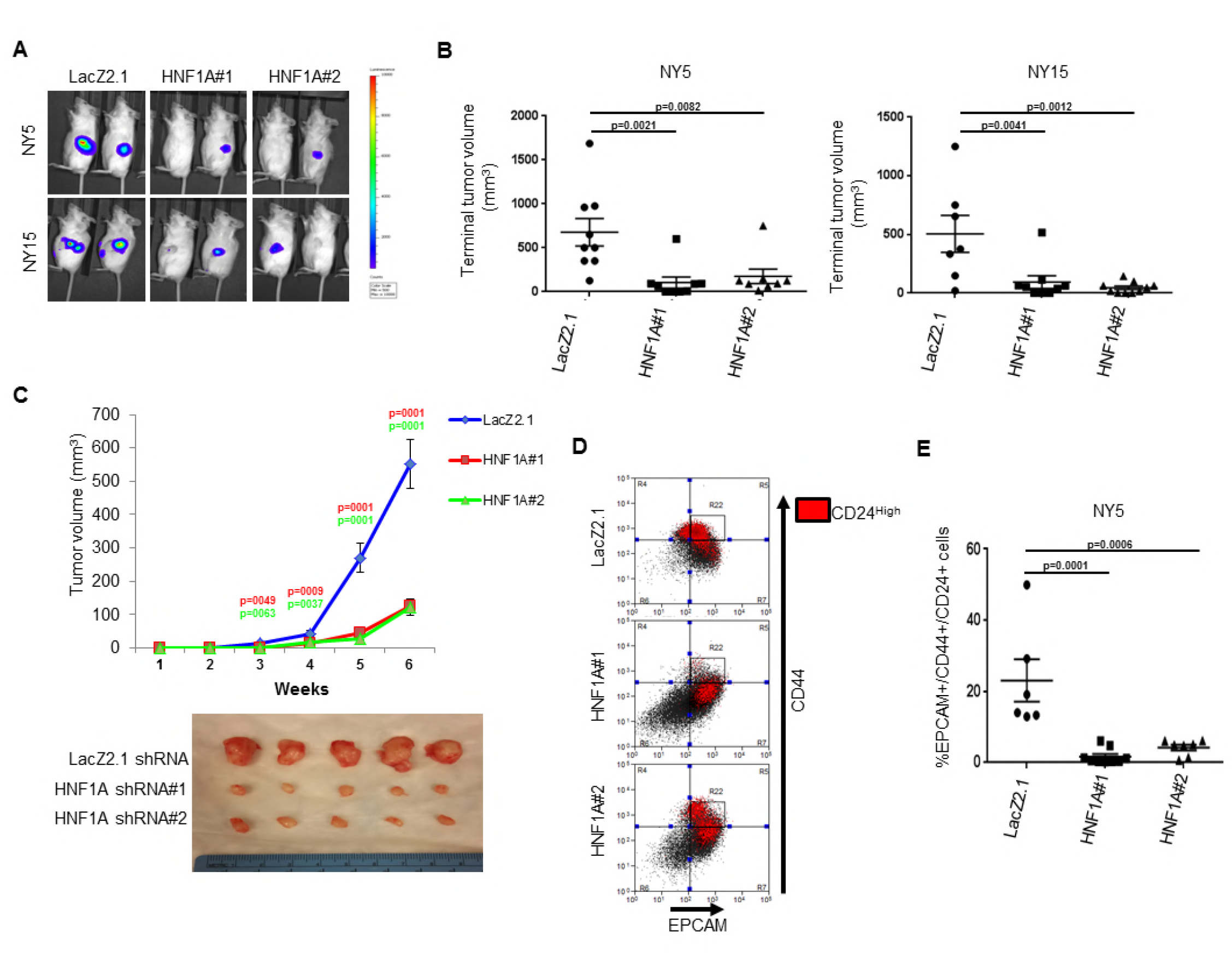
Knockdown of HNF1A impairs tumor growth and depletes CSCs in vivo. (A, B) 10,000 firefly luciferase-labeled NY5 and NY15 cells expressing control or HNF1A shRNAs were implanted orthotopically into the pancreata of NOD/SCID mice and monitored by IVIS imaging for 6 weeks (10 mice per group). Representative luminescence image of tumors prior to sacrifice is shown (A). Final tumor volumes determined during necropsy are quantitated in (B). Statistical difference was determined by oneway ANOVA with Dunnett’s multiple comparisons test. (C) 10^3^ control or HNF1A-depleted NY15 cells were implanted subcutaneously in NOD/SCID mice (10 mice per shRNA/bilateral injections) for 6 weeks. Tumors were measured by caliper to determine tumor growth (C, upper panel). Statistical difference was determined by one-way ANOVA with Dunnett’s multiple comparisons test. Red and green p values indicate LacZ2.1 vs. HNF1A#1 or #2, respectively. Representative tumors excised at sacrifice are shown (C, lower panel). (D, E) NY5 tumors from (A) were dissociated and stained for EPCAM, CD44, and CD24. Representative flow cytometry plots for recovered tumor cells are shown in (D), where the R22 gate denotes EPCAM^High^/CD44^High^ cells, and CD24+ cells are donated in red. Quantitation of EPCAM+/CD44+/CD24+ cells is shown in (E), n=6 tumors each for shRNA. Statistical difference was determined by one-way ANOVA with Dunnett’s multiple comparisons test. Related data can be found in **Supplementary Figure 5**.

**Figure 6:**
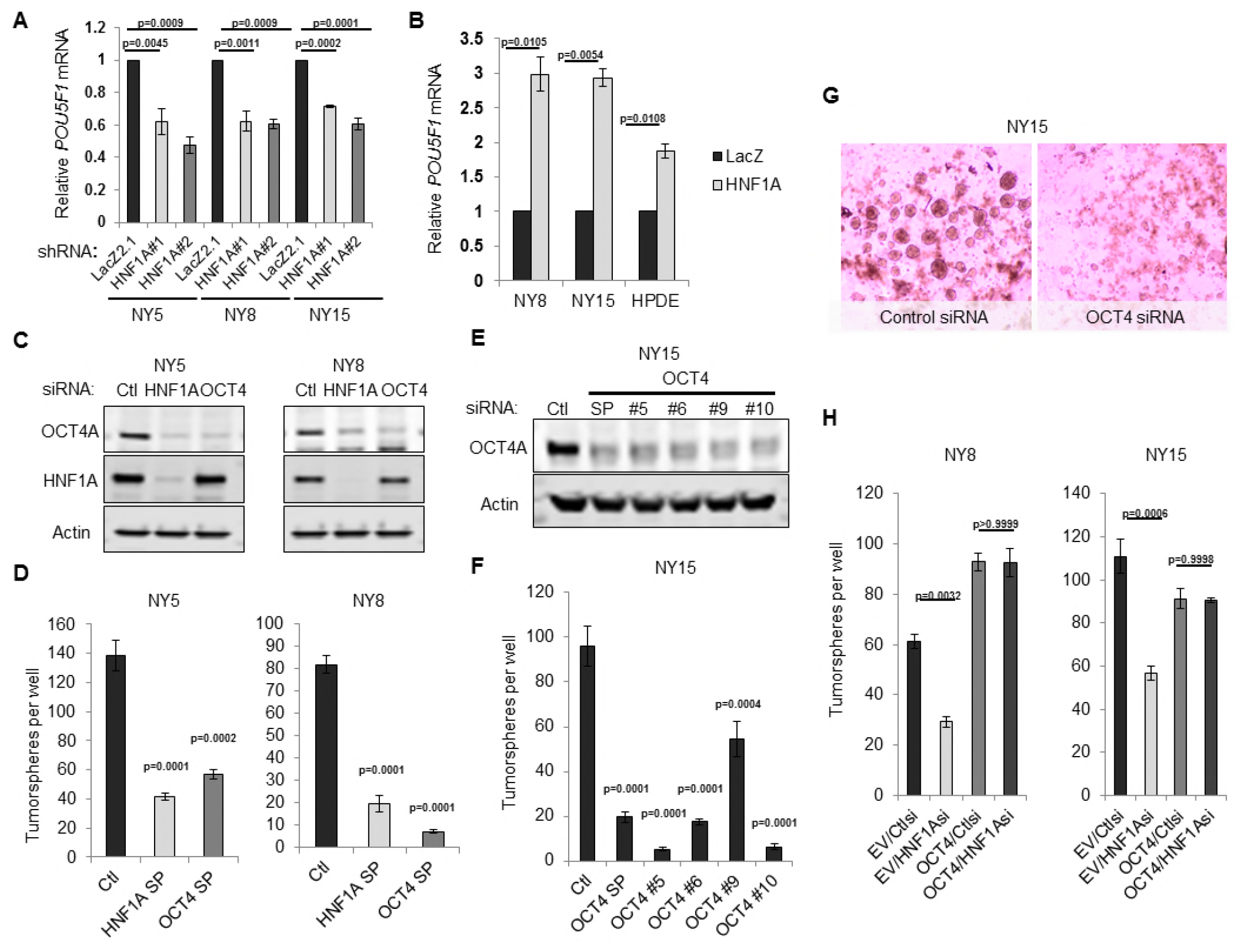
HNF1A regulates stemness through OCT4 regulation. (A) qRT-PCR analysis of *POU5F1* mRNA in NY5, NY8 and NY15 cells expressing control (LacZ2.1) or HNF1A shRNAs. *ACTB* was used as an internal control, n=3 biological replicates. Statistical difference was determined by one-way ANOVA with Dunnett’s multiple comparisons test. (B) LacZ or HNF1A was induced in NY8, NY15 or HPDE cells with doxycycline for 6 days. Levels of *POU5F1* mRNA were measured by qRT-PCR with *ACTB* as an internal control, n=3 biological replicates. Statistical difference was determined by unpaired t test with Welch’s correction. (C) Western blot of OCT4A and HNF1A protein in NY5 and NY8 cells transfected with OCT4 or HNF1A SMARTpool siRNA for 3 days. (D) NY5 and NY8 cells were transfected with OCT4 or HNF1A SMARTpool siRNA for 3 days and then grown in tumorsphere media on non-adherent plates (1500 cells/well). Spheres were quantitated 7 days later, n=3 biological replicates. Statistical difference was determined by one-way ANOVA with Dunnett’s multiple comparisons test. (E-G) NY15 cells were transfected with OCT4 SMARTpool (SP) siRNA or individual sequences for 3 days and either harvested to assess OCT4A knockdown by Western blot (E) or grown in tumorsphere media on non-adherent plates (1500 cells/well) (F). Spheres were quantitated 7 days later, n=3 biological replicates. Statistical difference was determined by one-way ANOVA with Dunnett’s multiple comparisons test. Representative spheres are shown in (G). (H) NY8 and NY15 cells transduced with OCT4A or empty vector control (EV) were transiently transfected with control (Ctl) or HNF1A-targeting siRNA for 72 hours, and then grown in tumorsphere media on non-adherent plates (1500 cells/well). Spheres were quantitated 7 days later, n=3 biological replicates. Statistical difference was determined by one-way ANOVA with Tukey’s multiple comparisons test. Related data can be found in **Supplementary Figure 6**.

To assess if the HNF1A-transcriptomic signature might serve as a prognostic tool for poor outcomes in pancreatic cancer patients, as has been observed with CSC signatures in other cancer types (Bartholdy et al., 2014, Eppert et al., 2011, Glinsky et al., 2005, Merlos-Suárez et al., 2011), we utilized The Cancer Genome Atlas (TCGA) dataset for PDA (PAAD cohort). Remarkably, 26/237 (11%) of HNF1A-activated genes and 17/137 (12.4%) of HNF1A-bound and -activate genes were significantly associated (log-rank test p-value <0.05) with poor survival outcome in patients with PDA (Figure 7D, E; Supplementary Figure 7A; Supplementary Table 2). By contrast, only 2/237 (0.8%) of HNF1A-activated genes and 1/137 (0.7%) of HNF1A-bound and activate genes were significantly associated with better survival outcomes. Importantly, expression of HNF1A-activated genes (both HNF1A-bound and unbound) was more likely to be associated with poor survival outcome in patients with PDA than genes selected at random (p<0.05), as determined by a permutation test (N=10,000) (see insets in Figure 7E; Supplementary Figure 6A). Conversely, expression of HNF1A-repressed genes was not predictive of survival (Supplementary Figure 6B, inset). Taken together, these data suggest that a HNF1A gene signature may predict poor outcomes in PDA.

**Figure 7:**
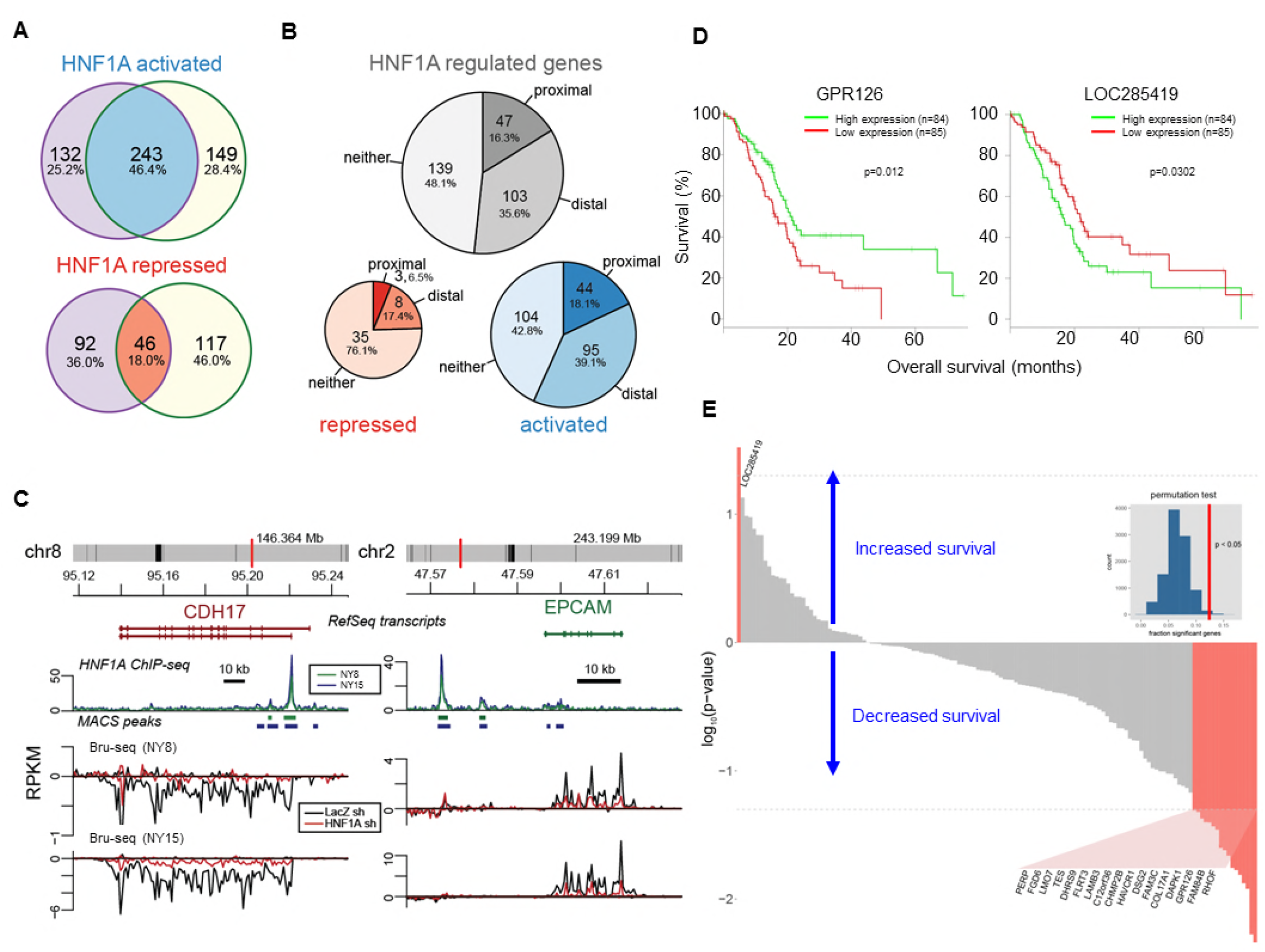
HNF1A regulates a transcriptional program associated with poor survival in PDA. (A) Venn diagrams illustrating overlapping genes with altered transcription (Bru-seq) following HNF1A knockdown in NY8 and NY15 cells. “HNF1A activated” genes denote genes that were downregulated by HNF1A shRNAs, while “HNF1A repressed” genes were upregulated by HNF1A shRNAs. Cells expressed shRNAs constitutively for >14 days prior to Bru-seq analysis. (B) Proportion of HNF1A shRNA-downregulated genes identified in both NY8 and NY 15 with HNF1A ChIP-seq peaks. Proximal peaks are +/-5 kb of the transcription start site (TSS) of a given gene and distal peaks are +/- 100 kb of a TSS. Peaks are recognized only if they are closer to the TSS of a given gene than to other expressed genes. (C) HNF1A ChIP-seq and HNF1A shRNA Bru-seq traces for the genes CDH17 and EPCAM in NY8 and NY15 cells. Traces represent normalized read coverage (in RPKM) across the indicated genomic ranges. MACS-identified ChIP peaks are represented by bars under the corresponding trace. (D) Representative Kaplan-Meier plot for direct HNF1A-target genes GPR126 and LOC285419. Red and green lines represent survival of the upper and lower 50^th^ percentile of *GPR126* and *LOC285419* mRNA expression in PDA patients, respectively. (E) HNF1A activated and bound genes were ranked according to log-rank test p-value and direction of survival based on TCGA PDA patient data. Y-axis units are log_10_-transformed p-values (higher magnitude indicates greater significance). Positive y-axis indicates association of gene expression with increased survival; negative y-axis indicates association of gene expression with reduced survival. Red bars indicate p-values < 0.05 (gray = not significant). Dotted line indicates p-value threshold of 0.05. Inset: histogram representing null distribution of a permutation test (N=10,000) for fraction of genes significantly associated with reduced survivability. Vertical red line represents value for set of HNF1A target genes. Related data can be found in **Supplementary Figure 7**.

## DISCUSSION

In this study, we identified the transcription factor HNF1A as putative regulator of a PCSC gene signature. Functional studies revealed that HNF1A was not only central to the regulation of this gene signature, but also PCSC function. Depletion of HNF1A effectively inhibited PDA cell growth, tumorsphere formation, and tumor growth, with a loss of PCSC numbers observed both *in vitro* and *in vivo*. Mechanistically, HNF1A appears to promote stemness through positive regulation of pluripotency factor POU5F1/OCT4. Finally, we found that expression of HNF1A-activated genes significantly predicted poor survival outcomes in patients with PDA. These data point to a novel oncogenic role for HNF1A in pancreatic cancer, particularly in PCSCs.

A clear role for HNF1A in PDA has not previously been established. An early study of the putative oncogene FGFR4, frequently expressed in PDA (Ohta et al., 1995), is directly regulated by HNF1A through intronic binding sites (Shah et al., 2002). More recently, 73% of PDA samples were found to stain positive for HNF1A (Kong et al., 2015). A more direct role for HNF1A in PDA has been suggested by multiple GWA studies implicating certain SNPs in HNF1A as risk factors for the development of PDA (Pierce and Ahsan, 2011a, Wei et al., 2012, Li et al., 2012). Nearly all of the identified HNF1A SNPs are non-coding and relatively common (minor allele frequencies between 30-40%), suggesting these SNPs may serve as potential contributing rather than driving factors in pancreatic tumorigenesis. Interestingly, PDA associated HNF1A SNPs rs7310409, rs1169300, and rs2464196 are also associated with both an elevated risk (1.5-2 fold) of developing lung cancer and elevated circulating C-reactive protein (CRP). A well-established direct target of HNF1A (Toniatti et al., 1990), CRP is downregulated in patients with inactivating mutations in HNF1A (Thanabalasingham et al., 2011). As several

PDA-associated SNPs are associated with elevated CRP, it is therefore possible that these SNPs augment the activity/expression of HNF1A rather than diminish it, as in the case of maturity-onset diabetes of the young 3 (MODY3) variants which reduce or abolish HNF1A expression or function. Still, a tumor suppressive role for HNF1A in PDA has also been proposed (Hoskins et al., 2014, Luo et al., 2015). In these studies HNF1A was found to possess pro-apoptotic/anti-proliferative properties contrary to the data in this study. Differences in these results may be technical in nature (control cells in Luo et al. exhibited unusually high baseline apoptosis approaching 50%), however it is also possible that the role of HNF1A may differ between different molecular subtypes of PDA (Bailey et al., 2016) or in a dynamic manner like fellow transcription factor PDX1 (Roy et al., 2016).

Our data on HPDE and HPNE cells support a partially transforming capacity for HNF1A, wherein it overcomes contact-inhibition and anchorage-dependent growth. As cooperation with oncogenic KRAS was observed in these cells, it is feasible that HNF1A provides additional oncogenic input, possibly by altering the differentiation state of KRAS-mutant, precancerous pancreatic cells or by expanding the resident stem cell/cancer stem cell population. Indeed, expression of HNF1A alone was sufficient to increase CD24 expression/positivity in both HPDE and HPNE cells.

Typically a marker of endodermal differentiation, HNF1A has not previously been reported as necessary for normal or cancer stem cells. HNF1A plays a critical role in the normal functionality of the endocrine pancreas, with hereditary inactivating mutations in the gene and promoter region resulting in MODY3, an autosomal dominant form of diabetes resulting from β cell insufficiency. Additionally, murine knockout models recapitulate the diabetic phenotype seen in humans (Lee et al., 1998), with elegant transcriptomic work demonstrating a requirement for murine Hnf1a in β cell proliferation (Servitja et al., 2009). The role for HNF1A in the exocrine pancreas is less clear, and compared to islet and liver cells in the latter study, we only identified 11 overlapping HNF1A-activated genes *(ANXA4, CEACAM1, CHKA, DPP4, HNF4A, HSD17B2, LGALS3, MTMR11, NR0B2, SLC16A5, TM4SF4)*, suggesting distinct activity for HNF1A in PDA compared to either β cells or the liver. Regulation of OCT4 by HNF1A is an especially exciting finding, connecting HNF1A with a previously unidentified role in regulating stemness. Although we have yet to determine precisely how HNF1A regulates *POU5F1* transcription, previously published HNF1A ChIP-seq data performed in HepG2 cells by The Encyclopedia of DNA Elements (ENCODE) project (Consortium, 2012), identified a region of enrichment of HNF1A upstream of the POU5F1/PSORS1C3 gene loci proximal to recently identified promoter regions for the genes (Malakootian et al., 2017), supporting direct regulation of OCT4 by HNF1A. Additionally, enrichment of this region by TATA-binding protein (TBP) and acetylated lysine 27 histone H3 supports the involvement of this region in the transcription of OCT4. As this region, rich in repetitive elements/retrotransposons, is not conserved between humans and rodents, it is possible the interaction between HNF1A and OCT4 is an acquisition of human evolution and may explain why OCT4 has not previously been identified as an HNF1A target. Our ChIP-seq data show a weak enrichment of HNF1A within this region and direct HNF1A ChIP shows a 2.5-4 fold enrichment in NY8 and NY15 cells. This lack of a robust identifiable interaction does not preclude direct regulation, but it suggests other indirect mechanisms may be just as important in regulation of OCT4 expression by HNF1A.

Given that 26 HNF1A-activated genes (17 direct targets) were found to be significantly associated with poor survival in patients with PDA, it is likely that multiple target genes contribute to HNF1A’s oncogenic influence, and future studies should be done to assess the functions of these genes in PDA to ascertain their value as either potential biomarkers or therapeutic targets. Further studies are also needed in regards to HNF1A’s role in the exocrine pancreas and whether its function is redirected during the development of PDA, particularly under the influence of oncogenic KRAS. Overall, this study further validates the importance of HNF1A to PDA while providing a novel and critical role for HNF1A in driving pancreatic cancer stem cells.

## MATERIALS AND METHODS

### Tumor growth assays

8-10 week old, evenly sex-mixed NOD/SCID mice were used for all experiments. Orthotopic implantation of PDA cells to the pancreas has previously been described (Abel et al., 2014). Briefly, mice were anesthetized with an intraperitoneal injection of 100 mg/kg ketamine/5 mg/kg xylazine, and a small left subcostal incision was performed. 10,000 PatGFP-Luc2-labeled tumor cells in a volume of 50 μl (1:1 volume of cell suspension in growth media and Matrigel) were injected into the tail of the pancreas using a 30-gauge needle. Weekly bioluminescent imaging of implanted orthotopic tumors in mice was performed using a Xenogen IVIS 200 Imaging System (Xenogen Biosciences, Cranbury, NJ). For subcutaneous implantation of tumor cells, 10,000 tumor cells in a volume of 50 μl (1:1 volume of cell suspension in growth media and Matrigel) was injected subcutaneously into both the left and right midflank regions of mice. Tumor growth was monitored weekly by digital caliper and tumor volumes calculated by the (length x width^2^)/2 method. All mice were sacrificed once any tumors reached 20mm^3^ in volume.

### Immunofluorescence and immunohistochemistry

Formalin-fixed, paraffin-embedded tumor samples were sectioned and processed for immunofluorescent staining by the University of Michigan ULAM Pathology Cores for Animal Research. Mason’s Trichrome staining was performed by the University of Michigan ULAM Pathology Cores for Animal Research. Immunohistochemistry was performed using a Ventana BenchMark Ultra autostainer. HNF1A antibody (GT4110) was used for immunohistochemistry at a 1:100 dilution. A PDA/normal pancreas tissue microarray was generated by the University of Michigan Department of Pathology.

### Microscopy

All microscopy was performed on an Olympus IX83 motorized inverted microscope with cellSens™ Dimension software (Olympus Corporation, Waltham, MA).

### Lentiviral constructs

Lentiviral destination vectors were generously provided by Dr. Andrew Aplin (Thomas Jefferson University). For construction of HNF1A, KRAS^G12D^, GFP and LacZ cDNA lentiviruses, pLentipuro3/TO/V5-DEST, pLentineo3/TO/V5-DEST, pLentihygro3/TO/V5-DEST were used. For OCT4A, pLenti6.3/UbC/V5-DEST was used. An EcoRV digested/re-ligated pLenti6.3/UbC/V5-DEST (removing the Gateway cloning element) was used as an empty vector control. For construction of shRNA lentiviruses, pLentipuro3/BLOCK-iT-DEST was used. Human HNF1A and KRAS^G12D^ were cloned from primary PDA cDNA into pENTR™/D-TOPO^®^ (Invitrogen). Human OCT4A was cloned from pCR4-TOPO clone BC117435 (Transomic Technologies) into pENTR™/D-TOPO^®^. LacZ and PatGFP (a variant of EGFP containing the following mutations: S31R, Y40N, S73A, F100S, N106T, Y146F, N150K, M154T, V164A, I168T, I172V, A207V) were also cloned into pENTR™/D-TOPO^®^ as control proteins. For labeling cells with firefly luciferase, PatGFP was fused to the N-terminus of firefly luciferase Luc2 (subcloned from pGL4.10) and cloned into pENTR/D-TOPO^®^ using Gibson Assembly (New England Biolabs). PatGFP-Luc2 was recombined into pLenti0.3/EF/V5-DEST, a modified version of pLenti6.3/UbC/V5-DEST with the human EF-1α promoter instead of the human UbC promoter and no downstream promoter/selective marker cassette, to generate pLenti0.3/EF/GW/PatGFP-Luc2. To generate the HNF1A-responsive reporter, the multiple cloning site and minimal promoter from pTA-Luc (Takara, Mountain View, CA) was subcloned upstream of PatGFP. 8 tandem repeats of the HNF1A-binding site with spacer nucleotides (CTTGGTTAATGATTAACCAGA) was cloned between the MluI and BglII sites of the multiple cloning site. LacZ2.1

(CACCAAATCGCTGATTTGTGTAGTCGTTCAAGAGACGACTACACAAATCAGCGA), HNF1A shRNA#1

(CACCGCTAGTGGAGGAGTGCAATTTCAAGAGAATTGCACTCCTCCACTAGC), and HNF1A shRNA#2

(CACCGTCCCTTAGTGACAGTGTCTATTCAAGAGATAGACACTGTCACTAAGGGAC) were cloned into pENTR™/H1/TO (Invitrogen). cDNA and shRNA constructs were recombined into their respective lentiviral plasmids using LR Clonase II (Invitrogen). The resulting constructs were packaged in 293FT cells as previously described.

### siRNA sequences

Non-targeting control (Cat#D-001810-01)

HNF1 A-targeting siRNA#1 (GGAGGAACCGTTTCAAGTG)

HNF1A-targeting siRNA#2 (GCAAAGAGGCACTGATCCA)

POU5F1-targeting siRNA#5 (CATCAAAGCTCTGCAGAAA)

POU5F1-targeting siRNA#6 (GATATACACAGGCCGATGT)

POU5F1-targeting siRNA#9 (GCGATCAAGCAGCGACTAT)

POU5F1-targeting siRNA#10 (TCCCATGCATTCAAACTGA)

### Cell culture

Low-passage xenograft tumors were cut into small pieces with scissors and then minced completely using sterile scalpel blades. Single cells were obtained described previously (Li et al., 2007). The cells were cultured in RPMI-1640 with GlutaMAX™-I supplemented with 10% FBS (Gibco), 1% antibiotic-antimycotic (Gibco), and 100μg/ml gentamicin (Gibco). The cells used in this article are passaged less than 10 times *in vitro*. HPDE cells were a generous gift from Dr. Craig Logsdon (MD Anderson) and were maintained in keratinocyte SFM supplemented (Invitrogen) with included EGF and bovine pituitary extract as well as 1% antibiotic-antimycotic and 100μg/ml gentamicin. All cells were routinely tested for mycoplasma contamination using the MycoScope PCR Detection kit (Genlantis, San Diego, CA) and only mycoplasma-free cells were used for experimentation.

### Soft agar assays

Low-melting agarose (Invitrogen) was dissolved in serum-free RPMI-1640 with GlutaMAX™-I to a final concentration of 2% at 60°C and cooled to 42°C. 200 μL per well 2% agarose was evenly spread at the bottom of a 24-well dish, followed by 250 μL of 0.6% agarose (diluted with complete keratinocyte SFM and supplemented with FBS to 2.5%), a 250 μL of 0.4% agarose/cell suspension, and a 250 μL of acellular 0.4% agarose. Each layer was allowed to solidify a 4°C for 10 minutes and then heated to 37°C prior to adding the next layer. 500ul of complete keratinocyte SFM and supplemented with 2.5% FBS was added atop each gel and replenished every 3 days.

### Flow cytometry

Flow cytometry was performed as described previously (Li et al., 2007). Cells were dissociated with 2.5% trypsin/EDTA solution, counted and transferred to 5 mL tubes, washed with HBSS supplemented with FBS twice and resuspended in HBSS/2% FBS at a concentration of 1 million cells/100 μL. Primary antibodies were diluted 1:40 in cell suspensions and incubated for 30 minutes on ice with occasional vortexing. Cells were washed twice with HBSS/2% FBS and incubated for 20 minutes on ice with APC-Cy7 Streptavidin diluted 1:200. Cells were washed twice with HBSS/2% FBS and resuspended in HBSS/2%FBS containing 3 μM 4’,6-diamidino-2-phenylindole (DAPI) (Invitrogen, Carlsbad, CA). Flow cytometry and sorting was done using a FACSAria (BD Biosciences, Franklin Lakes, NJ). Side scatter and forward scatter profiles were used to eliminate cell doublets, APC-Cy7 was used to exclude mouse cells. For PatGFP-Luc2 labeling, GFP+/DAPI-cells were isolated by sorting and expanded for one passage prior to implantation. For analysis of apoptosis, APC-conjugated Annexin V and Annexin V binding buffer (BD Biosciences) was used following manufacturer’s recommendations with 3 μM DAPI added immediately before analysis to stain permeable cells/necrotic debris.

### Microarray analysis

Flow sorted NY8 and NY15 P1, P2, and P3 cells were immediately used for RNA isolation using the RNeasy Plus Mini Kit coupled with RNase-free DNase set (Qiagen). Microarrays and analyses were performed by the University of Michigan DNA Sequencing Core. RNA labeling and hybridization was conducted using the Human Genome U133 Plus 2.0 microarray (Affymetrix, Santa Clara, CA). Probe signals were normalized and corrected according to background signal. Adjusted signal strength was used to generate quantitative raw values, which were log-transformed for all subsequent analyses.

### oPOSSUM 3.0 analysis of PCSC genes

Single Site Analysis (SSA) for human was used to detect over-represented conserved transcription factor binding sites in the 50 PCSC-enriched genes. The program was run using default settings, which included a conservation cutoff of 0.4, a matrix score threshold of 85%, and search region of 5,000 basepairs upstream and downstream of the start of transcription. The query was entered against a background of 24,752 genes in the oPOSSUM database.

### Quantitative reverse transcription-PCR (qRT-PCR)

Total RNA was extracted using RNeasy Plus Mini Kit coupled with RNase-free DNase set (Qiagen) and reverse transcribed with High Capacity RNA-to-cDNA Master Mix (Applied Biosystem). The resulting cDNAs were used for PCR using Power SYBR^®^ Green PCR Master Mix (Applied Biosystem) in triplicates. qPCR and data collection were performed on a ViiA™7 Real-Time PCR system (Invitrogen). Conditions used for qPCR were 95°C hold for 10 mins, 40 cycles of 95°C for 10 secs, 60°C for 15 secs, and 72°C for 20 secs. All quantitations were normalized to an endogenous control *ACTB*. The relative quantitation value for each target gene compared to the calibrator for that target is expressed as 2-(Ct-Cc) (Ct and Cc are the mean threshold cycle differences after normalizing to *ACTB)*.

### Tumorsphere cultures

Single cells were suspended in tumorsphere culture media containing 1% N2 supplement, 2% B27 supplement, 1% antibiotic-antimycotic, 20 ng/mL epidermal growth factor (Gibco, Carlsbad, CA), 20 ng/mL human bFGF-2 (Invitrogen), 10 ng/mL BMP4 (Sigma-Aldrich, St. Louis, MO), 10 ng/mL LIF (Sigma-Aldrich) and plated in 6-well Ultra-Low Attachment Plates (Corning, Corning, NY).

### siRNA transfection

siRNA were purchased from Dharmacon (Lafayette, CO) and were transfected at 25 nM using Lipofectamine^®^ RNAiMAX Reagent (Invitrogen). siRNA sequences can be found in the Supplementary Material and Methods.

### Western blotting

All lysates were boiled in 1x Laemmli sample buffer with β-mercaptoethanol for 5 minutes followed by electrophoresis on 4-20% Mini-PROTEAN TGX precast Tris-Glycine-SDS gels (Bio-Rad, Hercules, CA). Proteins were transferred to low-fluorescent PVDF (Bio-Rad) and incubated overnight in primary antibody at 1:1000 dilution. Blots were incubated in IRDye^®^-conjugated secondary antibodies at room temperature for 1 hour and imaged by Odyssey^®^ CLx imaging system (Li-Cor, Lincoln, NE). For western blotting, HNF1A (clone GT4110) and KRAS (ab55391) from Abcam (Cambridge, MA), β-Actin (clone AC-74) from Sigma-Aldrich, Cadherin-17 (CDH17) from Proteintech (Rosemont, IL), β-Galactosidase from Promega (Madison, WI) and RAS^G12D^, CD44, EPCAM, Cleaved Caspase-3 (D175), Cleaved Caspase-6 (D162), Cleaved Caspase-7 (D198), Cleaved Caspase-9 (D315), Cleaved Caspase-9 (D330), phospho-ERK1/2, phospho-AKT S473, OCT4A and GFP from Cell Signaling Technology (Danvers, MA). For flow cytometry, mouse anti-human EPCAM (CD326) clone HEA-125 was purchased from Miltenyi Biotec (San Diego, CA). Mouse anti-human CD44 clone G44-26, CD24 clone ML5, and CD26 (DPP4) clone M-A261 and APC-Cy7 Streptavidin were purchased from BD Biosciences (San Jose, CA). Biotinylated mouse anti-mouse H-2Kd/H-2Dd clone 34-1-2S was purchased from SouthernBiotech (Birmingham, AL).

### Chromatin immunoprecipitation sequencing (ChIP-seq)

A confluent 15cm culture plate of cells was used per immunoprecipitation. Cells were fixed with 1% formaldehyde for 10 minutes. Nuclei were collected and chromatin sheared to 1-10 nucleosomes using the SimpleChIP Plus Enzymatic Chromatin IP kit and protocol (Cell Signaling). HNF1A was immunoprecipitated with goat polyclonal antibody C-19 (Santa Cruz). Libraries from HNF1A-immunoprecipitated chromatin and input chromatin was prepared by the University of Michigan Sequencing Core and sequenced on the Illumina HiSeq 4000.

### Bromouridine labeling and sequencing (Bru-seq)

Nascent RNA labeling and sequencing (Bru-seq) was performed as previously described (Paulsen et al., 2013). For each shRNA (LacZ2.1, HNF1A shRNA#1, and #2), two replicates were performed in each cell line (NY8 and NY15). Cells were incubated in media containing 2 mM bromouridine (Bru) (Aldrich) for 30 minutes at 37°C. Total RNA was isolated after lysis in Trizol and Bru-RNA was isolated using anti-BrdU antibodies conjugated to magnetic beads. Strand-specific libraries were made using the Illumina TruSeq kit and sequenced on the Illumina HiSeq 4000 platform at the University of Michigan Sequencing Core (Ann Arbor, MI). Genes were recognized as differentially expressed in both cell lines if the fold change after knockdown was greater than 1.5 (and FDR <0.1 in NY15) and the mean RPKM for a given comparison was greater than 0.25 in either HNF1A shRNA#1 or shRNA#2 per cell line.

### Mapping and analysis of ChIP-seq and Bru-seq

For ChIP-seq, 52-base, single end reads were aligned to the human reference genome (hg19) using Bowtie v1.1.1 (with options: -n 3 -k 1 -m 1). Peaks were called using MACS v1.4.2 using the default options. MACS peaks overlapping ENCODE blacklist regions were removed (https://www.encodeproject.org/annotations/ENCSR636HFF). Each peak was assigned to the closest gene’s transcription start site (TSS). Then, for each TSS, the distance to the nearest peak was measured. If the nearest associated peak was within +/- 5 kb of the TSS, it was considered proximal. In the absence of a proximal peak, the nearest associated peak within +/- 100 kb of the TSS was considered distal. A gene was recognized as having a proximal or distal peak if at least one replicate in both cell lines identified a proximal or distal peak. If a gene was found to have both proximal and distal peaks (usually due to differences between replicates), the gene was identified as distal if it had distal peaks in both replicates of both cell lines, otherwise it was identified as neither. For Bru-seq, 52-base, single end reads were aligned first to ribosomal DNA (U13369.1) using Bowtie v0.12.8 and the remaining reads aligned to the human reference genome (hg19) using TopHat v1.4.1. Differential gene expression analysis was performed using DESeq v1.24.0 (R v3.3.1). ChIP-seq and Bru-seq data from this study are available at the NCBI Gene Expression Omnibus (GEO; accession # GSE108151).

### TCGA survival analysis

Gene expression and patient survival data for PAAD were obtained through the Broad Institute TCGA Genome Data Analysis Center (2016; Firehose stddata 2016_01_28 run; Broad Institute of MIT and Harvard; doi:10.7908/C11G0KM9). Clinical metadata were obtained from the Merge Clinical, Level 1 data set. Gene expression values were obtained from the Level 3 RSEM genes (normalized) data set and log_10_-transformed prior to analysis. Samples identified as tumors and of non-neuroendocrine origin were used. Genes were selected based on Bru-seq and/or ChIP-seq as per above and linked (where possible) to TCGA expression data through Entrez ID or HUGO gene symbol. Survival analysis, including Kaplan-Meier estimate and log-rank test, was performed using the R package survival (v2.40-1). Patients were stratified as “high” and “low” based on the median gene expression. For genes within a given set, the fraction of genes associated with reduced (or increased) survival with p < 0.05 was calculated. For each selected set of genes (e.g. HNF1A-activated genes), a permutation test (N=10,000) was performed using randomly-selected sets of genes expressed in NY8 and NY15 cells. The fraction of genes significantly associated with reduced (or increased) survival was calculated (as above) and the resulting null distribution used to test significance of the given selected set of genes. For N=10,000, the estimated error at p=0.05 is ±0.0034 (less than 10%).

### Other statistical analysis

The following methods are specific to analysis of the data represented in Figure 1–6 and Supplementary Figure 1, 3, and S5. Data are expressed as the mean ± SEM. Statistically significant differences between two groups was determined by the two-sided Student t-test for continuous data, while ANOVA was used for comparisons between multiple groups. Significance was defined as P < 0.05. GraphPad Prism 6 was used for these analyses.

### Study approval

All animal protocols were approved by University Committee for the Use and Care of Animals (UCUCA) at University of Michigan. Patient samples were collected under a protocol approved by the IRB at the The University of Michigan. All patients gave informed consent.

## AUTHOR CONTRIBUTIONS

Conceptualization, E.V.A., M.G. and D.M.S.; Methodology, E.V.A., M.G., B.M. and M.J.; Software, B.M.; Investigation, E.V.A., M.G., S.A., N.R., E.H., A.A.A., M.L.D., S.U., L.W., J.S., and H.C.C.; Resources, E.V.A, B.M., C.K., J.S., M.J. and D.M.S.; Validation, E.V.A., M.G., B. M. and C.K.; Formal Analysis, E.V.A., B.M. and C.K.; Writing – Original Draft, E.V.A., B.M. and D.M.S.; Data Curation, B.M.; Writing – Review & Editing, E.V.A., B.M., H.C.C. and D. M.S.; Visualization, E.V.A and B.M.; Supervision, M.W., H.C.C. and D.M.S., Project Administration, E.V.A, H.C.C. and D.M.S.; Funding Acquisition, E.V.A., H.C.C. and D.M.S.

## ACKNOWLEDGMENTS

We thank the University of Michigan Flow Cytometry Core facility for assistance with performing FACS analysis and sorting, the University of Michigan DNA Sequencing Core facility for assistance ChIP-seq and microarray setup and analysis, Armand Bankhead for bioinformatic consultation, and Michelle Paulsen for preparing samples for Bru-seq. The work was supported by the Pancreatic Cancer Action Network-AACR Pathway to Leadership Grant (16-70-25-ABEL) and the American Cancer Society Postdoctoral Fellowship (127662-PF-15-033-01-DDC) (to EVA) and the Gershenson Pancreatic Cancer Fund (DMS) and SKY Foundation (HC and DMS).

## SUPPLEMENTARY FIGURES AND FIGURE LEGENDS

**Supplementary Figure 1:**
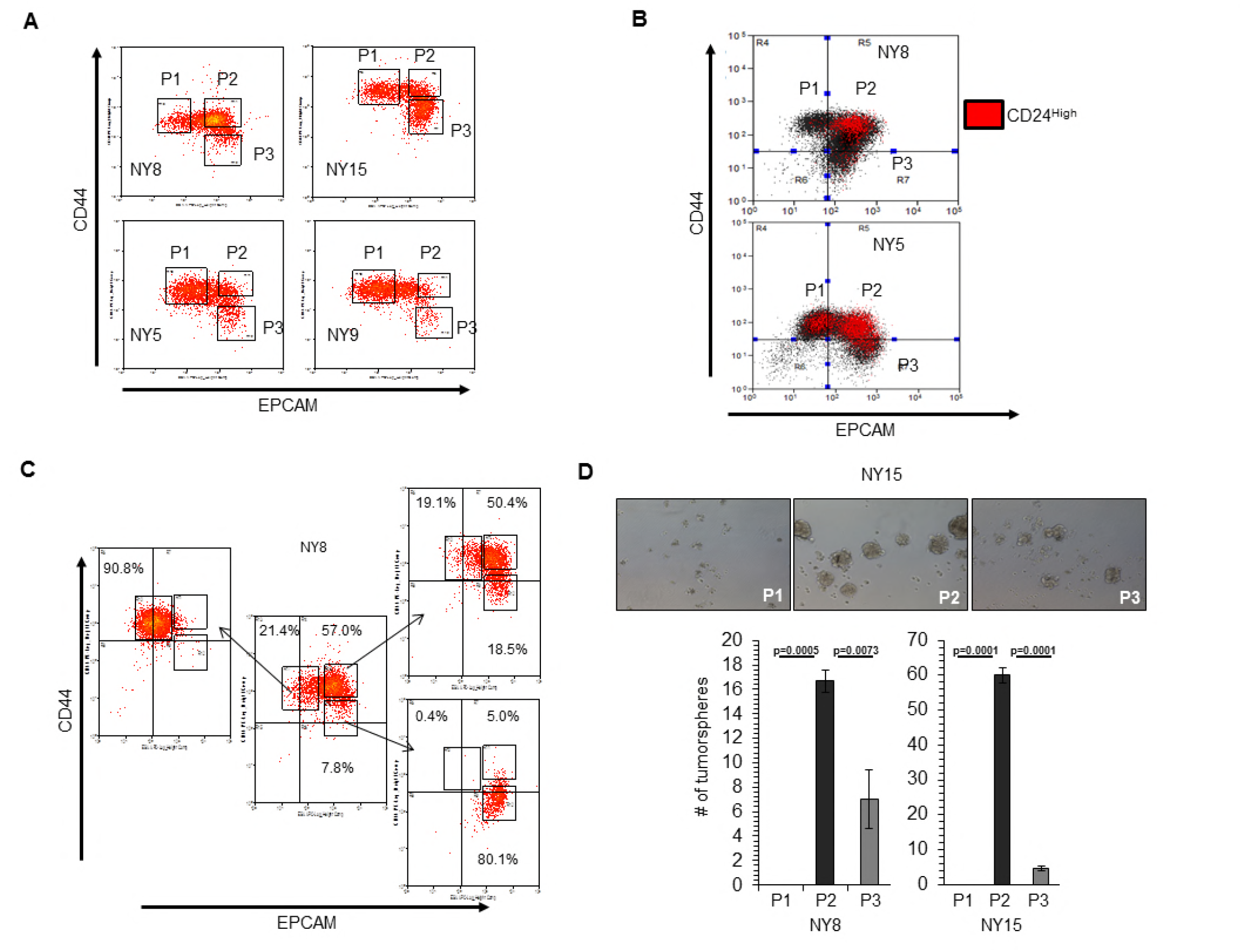
Cancer stem cell properties of PDA cell subpopulations. (A) Flow cytometry analysis of CD44 and EPCAM surface expression of 4 primary PDA samples. (B) Co-localization of CD24^High^ surface expression with EPCAM and CD44 expression is highlighted in red. (C) EPCAM^Low^/CD44^High^ (P1), EPCAM^High^/CD44^High^ (P2) and EPCAM^High^/CD44^Low^ (P3) NY8 cells were isolated by FACS and grown in culture for 17 days, followed by flow cytometry for analysis for CD44 and EPCAM expression. (D) Isolated subpopulations were grown in tumorsphere media on non-adherent plates (500 cells/well). The number of tumorspheres formed after 6 days were counted (n=3). Statistical difference was determined by one-way ANOVA with Tukey’s multiple comparisons test. Representative images of spheres (100X magnification) are shown in the right panel.

**Supplementary Figure 2:**
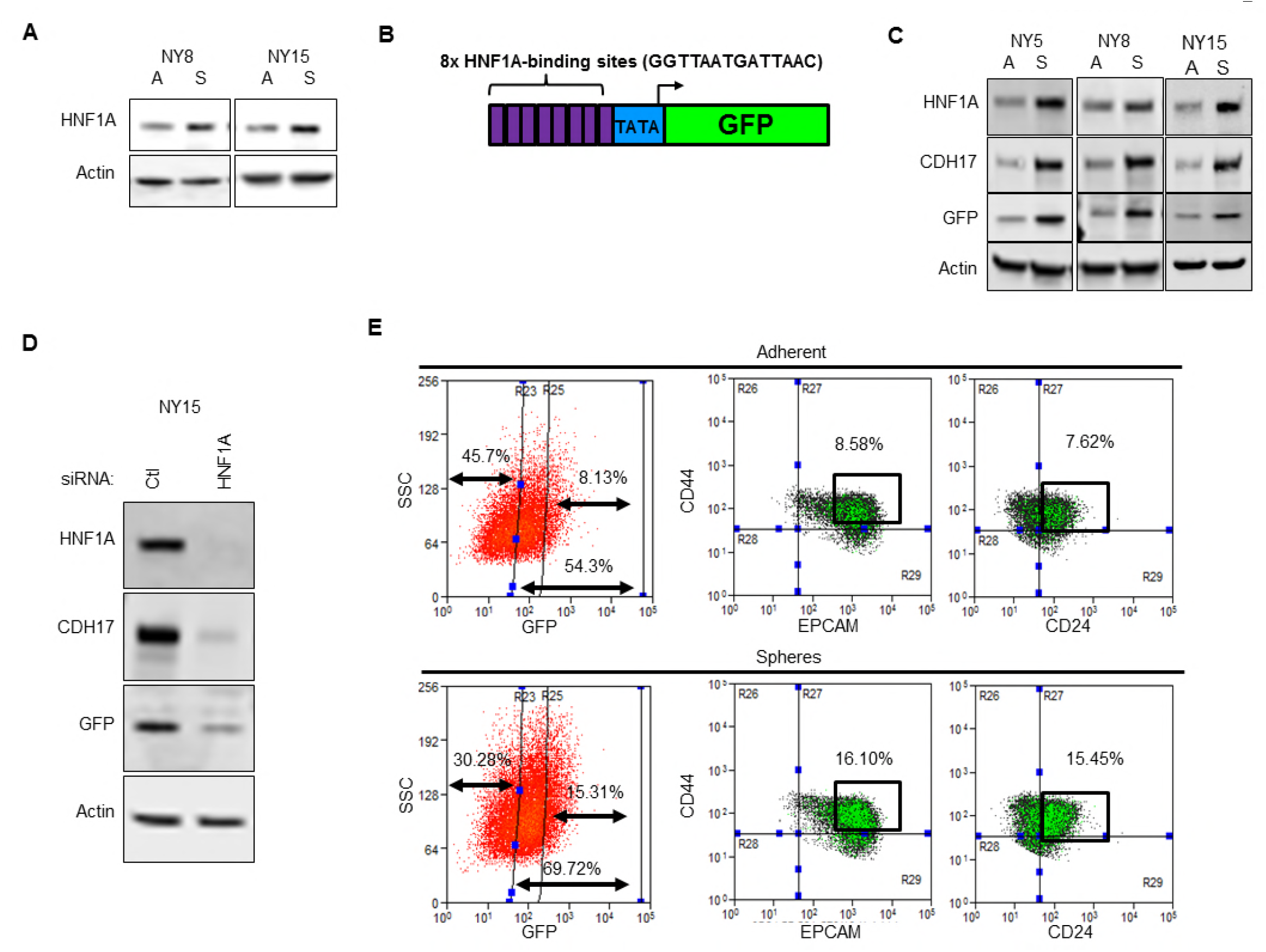
HNF1A is elevated in tumorspheres. (A) NY8 and NY15 cells were grown either in non-adherent conditions (labeled S) or under adherent conditions (labeled A) for 7 days. Lysates were collected and western blotted for HNF1A, CDH17, and Actin. (B) Schematic representation of HNF1A-responsive reporter construct showing 8 tandem copies of the HNF1A-binding site upstream of a minimal promoter (pTA) driving GFP expression. (C) HNF1A-reporter-labeled NY5, NY8, and NY15 cells were grown under adherent (A) or tumorsphere (S) conditions for 7 days, lysed and immunoblotted for HNF1A, CDH17, GFP, and Actin. (D) Reporter cells were transfected for 7 days with control (Ctl) or HNF1A-targeting siRNA, lysed and immunoblotted for HNF1A, CDH17, GFP, and Actin. (E) Reporter cells grown under adherent or tumorsphere conditions were collected by trypsinization and analyzed by flow cytometry for GFP fluorescence and EPCAM, CD44, and CD24 surface staining. The percentage of EPCAM+/CD44+/GFP+ or CD44+/CD24+/GFP+ cells is indicated.

**Supplementary Figure 3:**
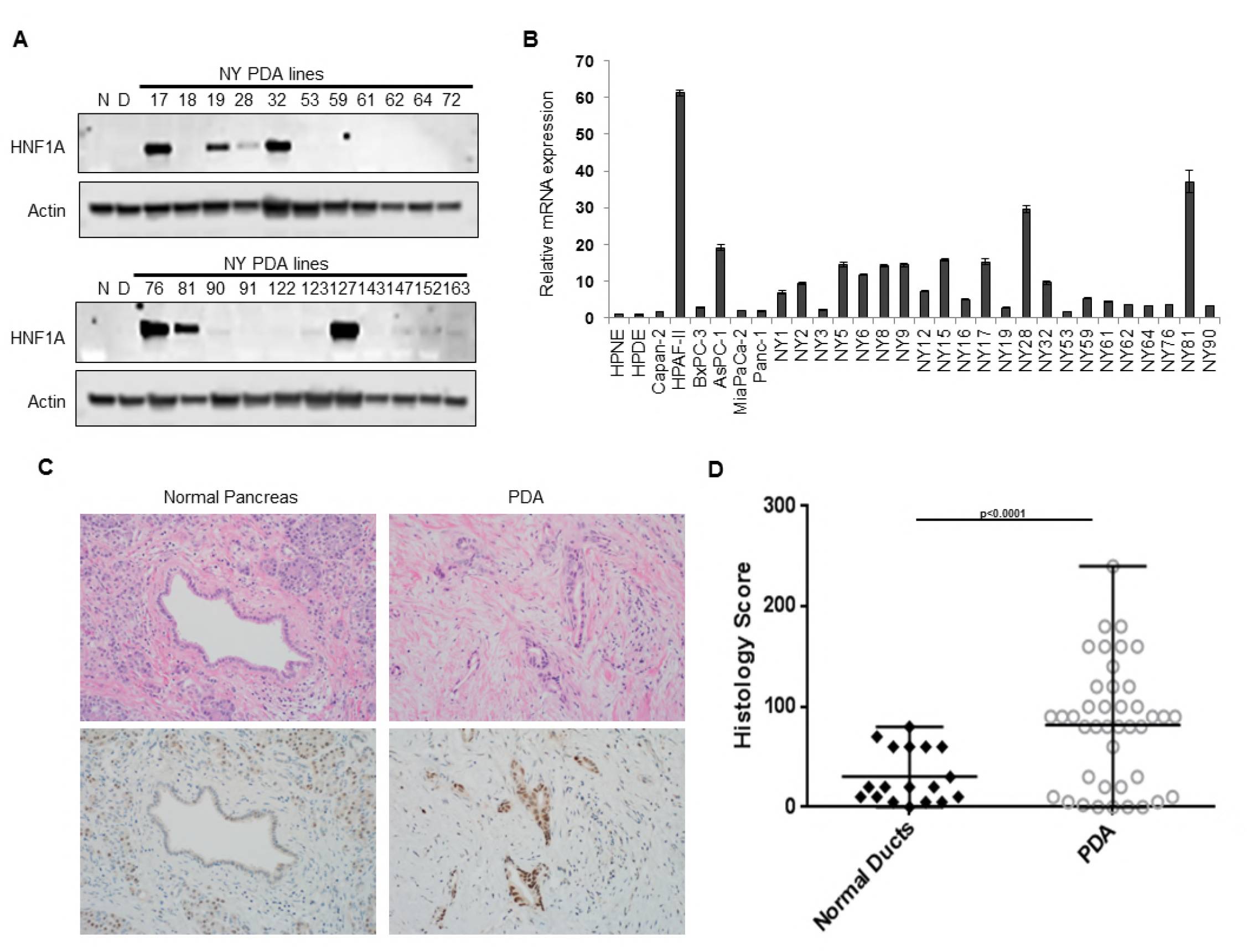
HNF1A expression in PDA cells and patient samples. (A) Western blot analysis of HNF1A expression in a panel of primary PDA lines compared to immortalized pancreatic ductal cell line HPNE (N) and HPDE (D). (B) qRT-PCR analysis of HNF1A expression in a panel of conventional and primary PDA lines. All qRT-PCR was normalized to an *ACTB* internal control, (n=3). (C, D) A pancreatic cancer tissue microarray (TMA) consisting of 41 PDA samples and 18 normal pancreas controls was immunostained for HNF1A. Representative hematoxylin and eosin staining and HNF1A staining of normal pancreas and PDA are shown in (C). Histology scores (HNF1A staining intensity x % HNF1A positive cells) for normal pancreatic ducts (n=18) versus PDA (n=41) is shown in (D). Statistical difference was determined by unpaired t test with Welch’s correction.

**Supplementary Figure 4:**
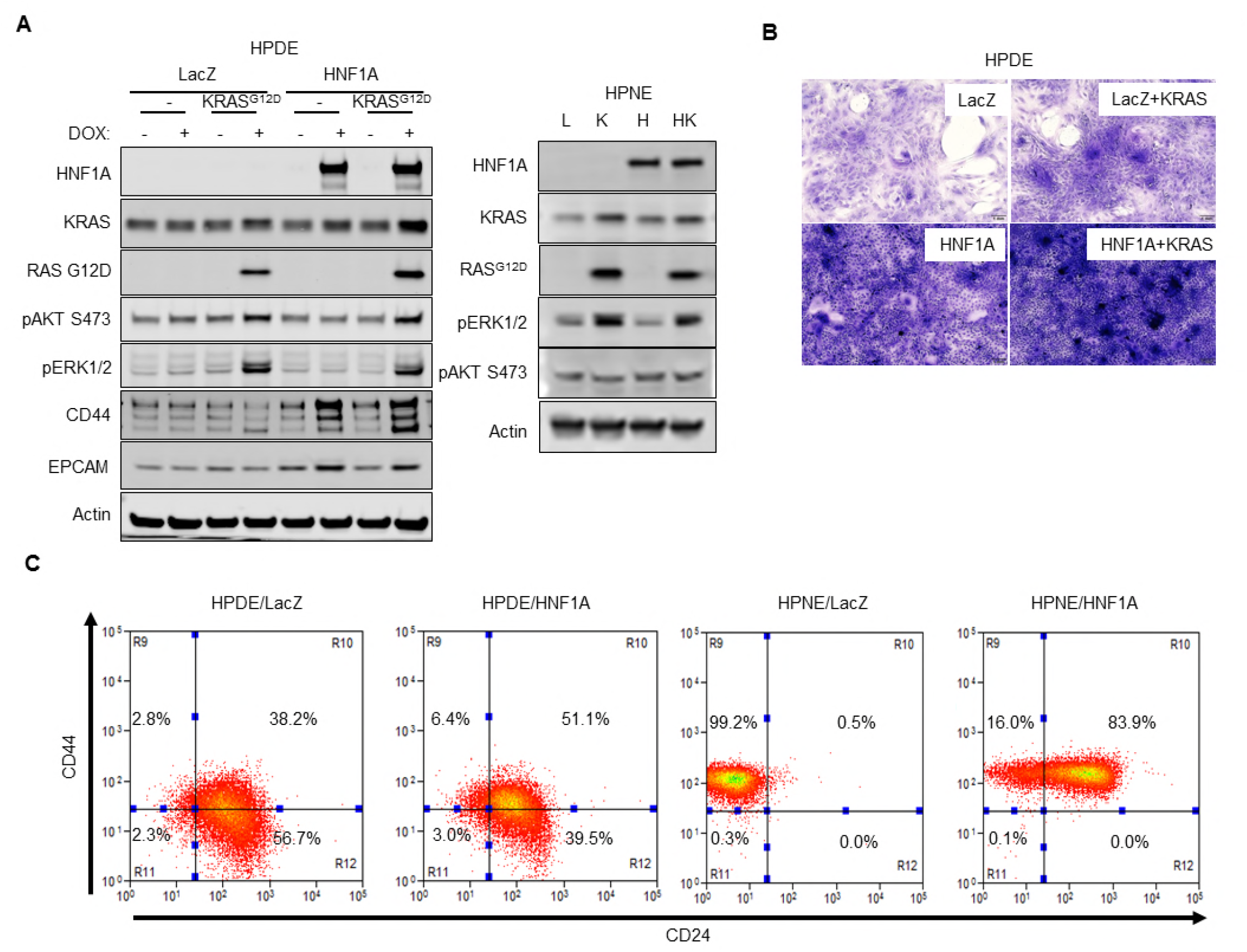
Overexpression of HNF1A and mutant KRAS in HPDE and HPNE cell. (A) HPDE and HPNE cells expressing LacZ (L), KRAS^G12D^ (K), HNF1A (H), or HNF1A with KRAS^G12D^ (HK) were analyzed by Western blot to confirm transgene expression and activation of downstream signaling events. For doxycycline-inducible HPDE cells, cells were treated ± doxycycline (Dox) (100 ng/ml) for 7 days prior to analysis. (B) HPDE cells expressing inducible LacZ, HNF1A, LacZ with KRAS^G12D^, or HNF1A with KRAS^G12D^ were grown to super-confluency (2 weeks) in the presence of 10% FBS + Dox to test for changes in contact-inhibition. Cells were fixed and stained with crystal violet. Representative images (100X magnification) are shown. (C) Flow cytometry analysis of CD44 and CD24 surface expression on HPDE cells expressing LacZ or HNF1A for 7 days or HPNE cells expressing LacZ or HNF1A.

**Supplementary Figure 5:**
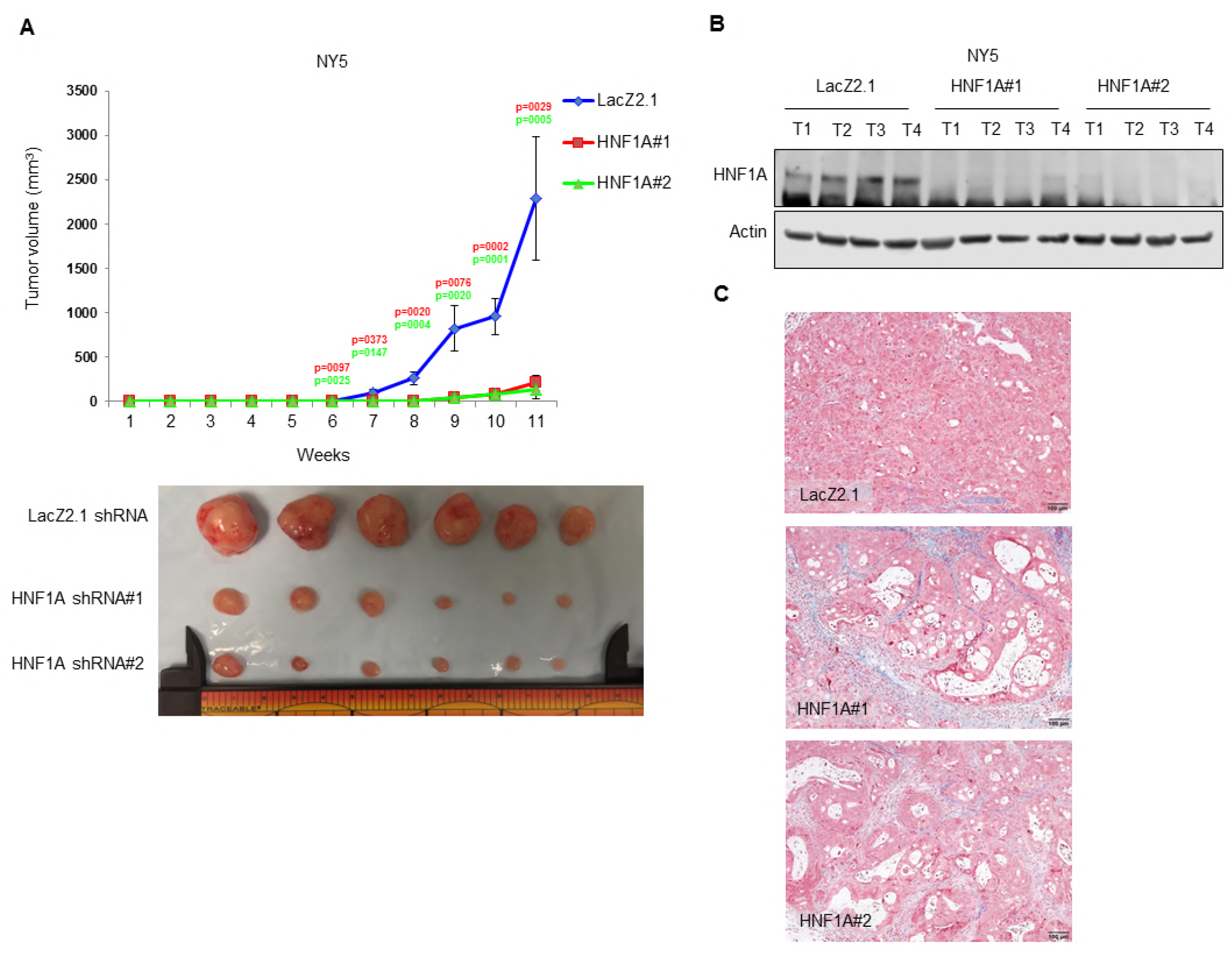
Effects of HNF1A depletion on PDA xenograft biology. (A) 10^3^ control or HNFlA-depleted NY5 cells were implanted subcutaneously in NOD/SCID mice (10 mice per shRNA/bilateral injections) for 6 weeks. Tumors were measured by caliper to determine tumor growth. Statistical difference was determined by one-way ANOVA with Dunnett’s multiple comparisons test. Red and green p values indicate LacZ2.1 vs. HNF1A#1 or #2, respectively. Representative tumors are shown (lower panel). (B) Western blot analysis of HNF1A and Actin from NY5 tumors expressing LacZ2.1 or HNF1A shRNAs (4 tumors per group) demonstrating persistent knockdown *in vivo*. (C) Representative Mason’s Trichrome stained NY5 tumor sections from LacZ2.1 or HNF1A shRNA-expressing tumors.

**Supplementary Figure 6:**
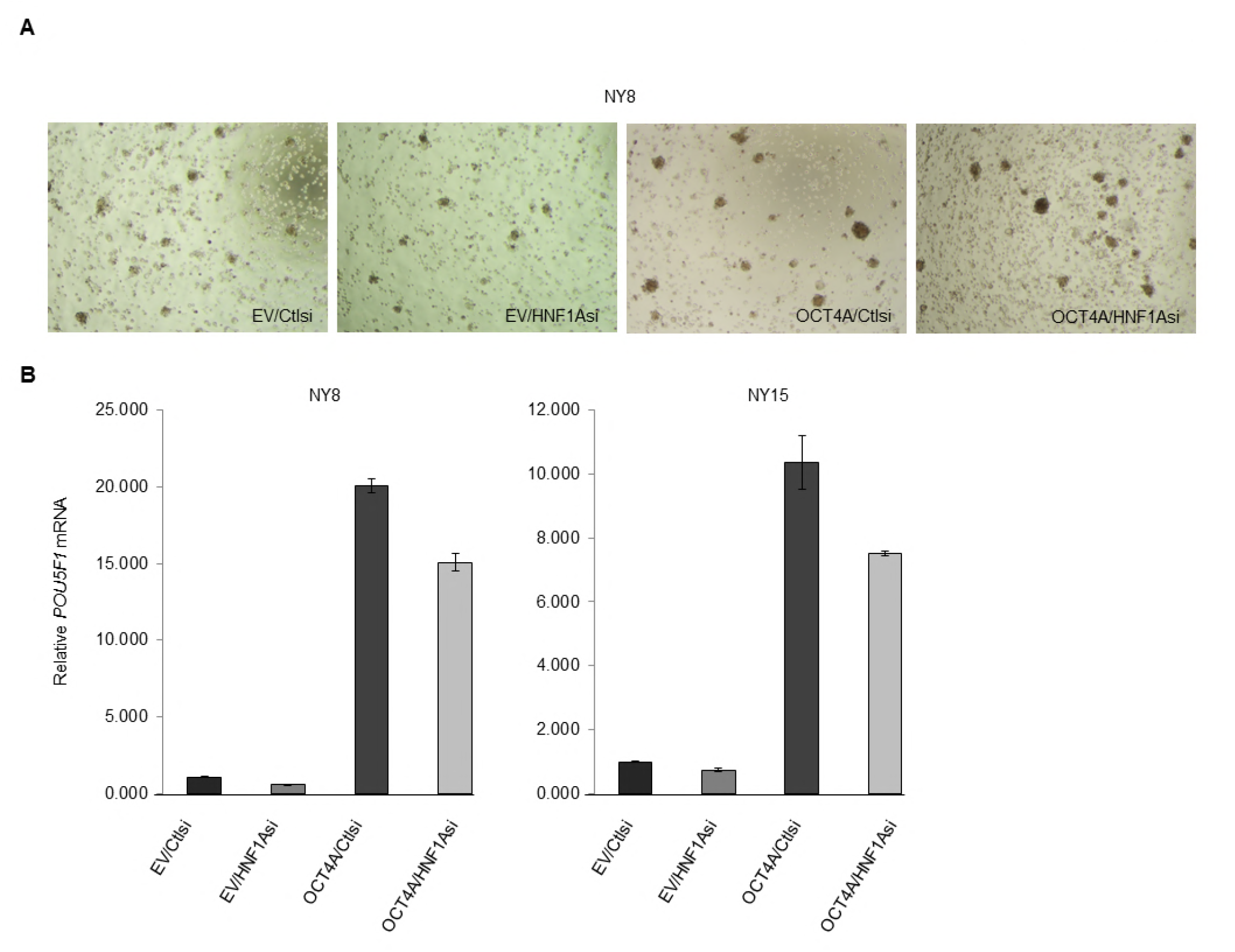
Rescue of OCT4 expression in PDA cells. (A) NY8 cells transduced with OCT4A or empty vector control (EV) were transiently transfected with control (Ctl) or HNF1A-targeting siRNA for 72 hours, and then grown in tumorsphere media on non-adherent plates (1500 cells/well). Representative images of tumorspheres are shown. (B) Quantitative RT-PCR for OCT4 transcript *(POU5F1)* was measured in NY8 and NY15 EV or OCT4 transduced cells transfected with control or HNF1A siRNA for 72 hours. *ACTB* transcript was used as an internal control, n=3.

**Supplementary Figure 7:**
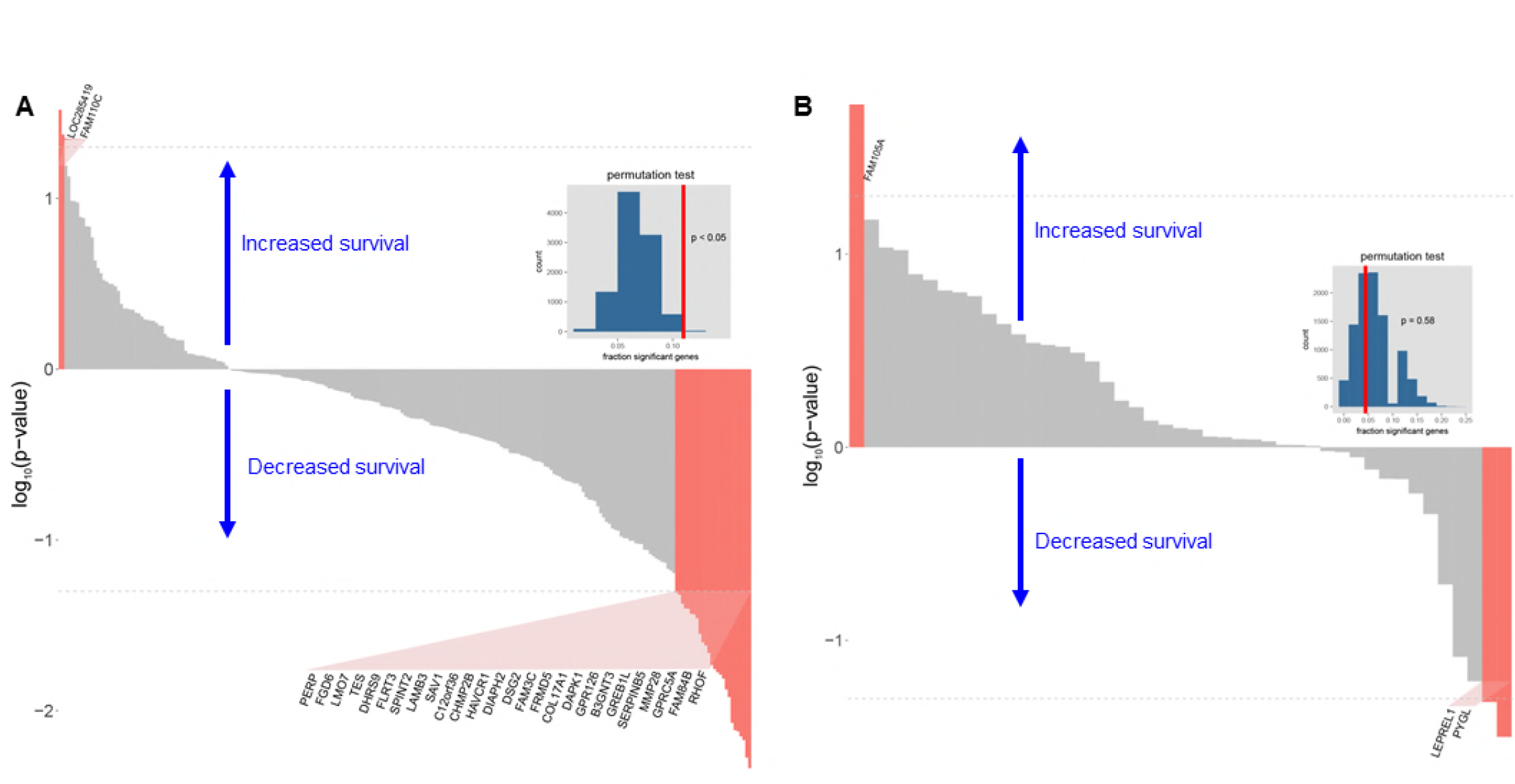
Association of HNF1A-responsive genes and survival in PDA. (A, B) HNF1A-responsive genes were ranked according to log-rank test p-value and direction of survivability based on TCGA PDA patient data. All HNF1A activated genes (bound and unbound) (A) and repressed genes (B) were analyzed. Y-axis units are log_10_-transformed p-values (higher magnitude indicates greater significance). Positive y-axis indicates association of gene expression with increased survival; negative y-axis indicates association of gene expression with reduced survival. Red bars indicate p < 0.05 (gray = not significant). Dotted line indicates p-value threshold of 0.05. Inset: histogram representing null distribution for fraction of genes significantly associated with increased survivability. Vertical red lines represents value for set of HNF1A-activated or -repressed genes, respectively. For N=10,000, the estimated error at p=0.05 is ±0.0034 (less than 10%).

## SUPPLEMENTARY TABLES

**Supplementary Table 1:** Cancer stem cell frequencies in PDA subpopulations. Limiting dilution assay was performed with sorted NY15 cells injected subcutaneously in NOD/SCID mice. The resultant numbers of tumors/injection is tabulated with estimated cancer stem cell frequencies calculated by extreme limiting dilution analysis (ELDA).

| Sub-population (NY15) | 2000 cells/ animal | 200 cells/ animal | 20 cells/ animal | Estimated CSC frequency (lower-upper confidence intervals) | Chisq (vs. HH) | DF | Pr(>Chisq) (vs. HH) |
| --- | --- | --- | --- | --- | --- | --- | --- |
| P1 | 1/5 | 0/5 | 0/5 | 1 in 10067 (71034-1427) | 12.1 | 1 | 0.000492 |
| P2 | 5/5 | 2/5 | 0/5 | 1 in 406 (1250-406) | N/A | N/A | N/A |
| P3 | 3/5 | 1/5 | 0/5 | 1 in 1869 (5288-1869) | 4.12 | 1 | 0.0424 |

**Supplementary Table 2: Data for generating PDA subpopulation heatmap and HNF1A target gene data (Excel spreadsheet)**

## REFERENCES

Abel, E. V., Kim, E. J., Wu, J., Hynes, M., Bednar, F., Proctor, E., Wang, L., Dziubinski, M. L. & Simeone, D. M. 2014. The Notch Pathway Is Important in Maintaining the Cancer Stem Cell Population in Pancreatic Cancer. PLoS ONE, 9, e91983.

Babajko, S., Tronche, F. & Groyer, A. 1993. Liver-specific expression of human insulin-like growth factor binding protein 1: functional role of transcription factor HNF1 in vivo. Proc Natl Acad Sci U S A, 90, 272–276.

Bailey, P., Chang, D. K., Nones, K., Johns, A. L., Patch, A.-M., Gingras, M.-C., Miller, D. K., Christ, A. N., Bruxner, T. J. C., Quinn, M. C., Nourse, C., Murtaugh, L. C., Harliwong, I., Idrisoglu, S., Manning, S., Nourbakhsh, E., Wani, S., Fink, L., Holmes, O., Chin, V., Anderson, M. J., Kazakoff, S., Leonard, C., Newell, F., Waddell, N., Wood, S., Xu, Q., Wilson, P. J., Cloonan, N., Kassahn, K. S., Taylor, D., Quek, K., Robertson, A., Pantano, L., Mincarelli, L., Sanchez, L. N., Evers, L., Wu, J., Pinese, M., Cowley, M. J., Jones, M. D., Colvin, E. K., Nagrial, A. M., Humphrey, E. S., Chantrill, L. A., Mawson, A., Humphris J., Chou A., Pajic, M., Scarlett, C. J., Pinho, A. V., Giry-Laterriere M., Rooman, I., Samra, J. S., Kench, J. G., Lovell, J. A., Merrett, N. D., Toon, C. W., Epari, K., Nguyen, N. Q., Barbour, A., Zeps, N., Moran-Jones K., Jamieson, N. B., Graham, J. S., Duthie, F., Oien, K., Hair, J., Grützmann, R., Maitra, A., Iacobuzio-DonahueC. A., Wolfgang, C. L., Morgan, R. A., Lawlor, R. T., Corbo, V., Bassi, C., Rusev, B., Capelli, P., Salvia, R., Tortora, G., Mukhopadhyay, D., Petersen, G. M., AUSTRALIAN PANCREATIC CANCER GENOME, I., Munzy, D. M., Fisher, W. E., Karim, S. A., Eshleman, J. R., Hruban, R. H., Pilarsky, C., Morton, J. P., Sansom, O. J., Scarpa, A., Musgrove, E. A., Bailey, U.-M. H., Hofmann, O., Sutherland, R. L., Wheeler, D. A., Gill, A. J., Gibbs, R. A., Pearson, J. V., et al. 2016. Genomic analyses identify molecular subtypes of pancreatic cancer. Nature, 531, 47–52.

Bartholdy, B., Christopeit, M., Will, B., Mo, Y., Barreyro, L., Yu, Y., Bhagat, T. D., Okoye-Okafor U. C., Todorova, T. I., Greally, J. M., Levine, R. L., Melnick, A., Verma, A. & Steidl, U. 2014. HSC commitment–associated epigenetic signature is prognostic in acute myeloid leukemia. J Clin Invest, 124, 1158–1167.

Boj, S. F., Párrizas, M., Maestro, M. A. & Ferrer, J. 2001. A transcription factor regulatory circuit in differentiated pancreatic cells. Proc Natl Acad Sci U.S.A., 98, 14481–14486.

Bonnet, D. & Dick, J. E. 1997. Human acute myeloid leukemia is organized as a hierarchy that originates from a primitive hematopoietic cell. Nat. Med., 3, 730–737.

Collisson, E. A., Sadanandam, A., Olson, P., Gibb, W. J., Truitt, M., Gu, S., Cooc, J., Weinkle, J., Kim, G. E., Jakkula, L., Feiler, H. S., Ko, A. H., Olshen, A. B., Danenberg, K. L., Tempero, M. A., Spellman, P. T., Hanahan, D. & Gray, J. W. 2011. Subtypes of pancreatic ductal adenocarcinoma and their differing responses to therapy. Nat Med, 17, 500–503.

Consortium, E. P. 2012. An integrated encyclopedia of DNA elements in the human genome. Nature, 489, 57–74.

Eppert, K., Takenaka, K., Lechman, E. R., Waldron, L., Nilsson, B., Van Galen, P., Metzeler, K. H., Poeppl, A., Ling, V., Beyene, J., Canty, A. J., Danska, J. S., Bohlander, S. K., Buske, C., Minden, M. D., Golub, T. R., Jurisica, I., Ebert, B. L. & Dick, J. E. 2011. Stem cell gene expression programs influence clinical outcome in human leukemia. Nat. Med., 17, 1086–1093.

Glinsky, G. V., Berezovska, O. & Glinskii, A. B. 2005. Microarray analysis identifies a death-from-cancer signature predicting therapy failure in patients with multiple types of cancer. J Clin Invest, 115, 1503–1521.

Graham, S. M., Jørgensen, H. G., Allan, E., Pearson, C., Alcorn, M. J., Richmond, L. & Holyoake, T. L. 2002. Primitive, quiescent, Philadelphia-positive stem cells from patients with chronic myeloid leukemia are insensitive to STI571 in vitro. Blood, 99, 319–325.

Gu, N., Tsuda, M., Matsunaga, T., Adachi, T., Yasuda, K., Ishihara, A. & Tsuda, K. 2008. Glucose regulation of dipeptidyl peptidase IV gene expression is mediated by hepatocyte nuclear factor-1alpha in epithelial intestinal cells. Clin Exp Pharmacol Physiol, 35, 1433–9.

Hermann, P. C., Huber, S. L., Herrler, T., Aicher, A., Ellwart, J. W., Guba, M., Bruns, C. J. & Heeschen, C. 2007. Distinct populations of cancer stem cells determine tumor growth and metastatic activity in human pancreatic cancer. Cell Stem Cell, 1, 313–323.

Herreros-Villanueva M., Zhang, J. S., Koenig, A., Abel, E. V., Smyrk, T. C., Bamlet, W. R., De Narvajas, A. A. M., Gomez, T. S., Simeone, D. M., Bujanda, L. & Billadeau, D. D. 2013. SOX2 promotes dedifferentiation and imparts stem cell-like features to pancreatic cancer cells. Oncogenesis, 2, e61.

Hoskins, J. W., Jia, J., Flandez, M., Parikh, H., Xiao, W., Collins, I., Emmanuel, M. A., Ibrahim, A., Powell, J., Zhang, L., Malats, N., Bamlet, W. R., Petersen, G. M., Real, F. X. & Amundadottir, L. T. 2014. Transcriptome analysis of pancreatic cancer reveals a tumor suppressor function for HNF1A. Carcinogenesis, 35, 2670–2678.

Kim, M. P., Fleming, J. B., Wang, H., Abbruzzese, J. L., Choi, W., Kopetz, S., Mcconkey, D. J., Evans, D. B. & Gallick, G. E. 2011. ALDH activity selectively defines an enhanced tumor-initiating cell population relative to CD133 expression in human pancreatic adenocarcinoma. PLoS ONE, 6, e20636.

Kong, B., Wu, W., Valkovska, N., Jäger, C., Hong, X., Nitsche, U., Friess, H., Esposito, I., Erkan, M., Kleeff, J. & Michalski, C. W. 2015. A common genetic variation of melanoma inhibitory activity-2 labels a subtype of pancreatic adenocarcinoma with high endoplasmic reticulum stress levels. Sci Rep, 5, 8109.

Kwon, A. T., Arenillas, D. J., Hunt, R. W. & Wasserman, W. W. 2012. oPOSSUM-3: Advanced Analysis of Regulatory Motif Over-Representation Across Genes or ChIP-Seq Datasets. G3, 2, 987–1002.

Lee, J., Kim, H. K., Rho, J.-Y., Han, Y.-M. & Kim, J. 2006. The Human OCT-4 Isoforms Differ in Their Ability to Confer Self-renewal. J Biol Chem, 281, 33554–33565.

Lee, Y.-H., Sauer, B. & Gonzalez, F. J. 1998. Laron Dwarfism and Non-Insulin-Dependent Diabetes Mellitus in the Hnf-1α Knockout Mouse. Mol. Cell. Biol., 18, 3059–3068.

Li, C., Heidt, D. G., Dalerba, P., Burant, C. F., Zhang, L., Adsay, V., Wicha, M., Clarke, M. F. & Simeone, D. M. 2007. Identification of pancreatic cancer stem cells. Cancer Research, 67, 1030–1037.

Li, C., Wu, J. J., Hynes, M., Dosch, J., Sarkar, B., Welling, T. H., Pasca Di Magliano, M. & Simeone, D. M. 2011. c-Met is a marker of pancreatic cancer stem cells and therapeutic target. Gastroenterology, 141, 2218–2227.e5.

Li, D., Duell, E. J., Yu, K., Risch, H. A., Olson, S. H., Kooperberg, C., Wolpin, B. M., Jiao, L., Dong, X., Wheeler, B., Arslan, A. A., Bueno-De-Mesquita, H. B., Fuchs, C. S., Gallinger, S., Gross, M., Hartge, P., Hoover, R. N., Holly, E. A., Jacobs, E. J., Klein, A. P., Lacroix, A., Mandelson, M. T., Petersen, G., Zheng, W., Agalliu, I., Albanes, D., Boutron-Ruault, M.-C., Bracci, P. M., Buring, J. E., Canzian, F., Chang, K., Chanock, S. J., Cotterchio, M., Gaziano, J. M., Giovannucci, E. L., Goggins, M., Hallmans, G., Hankinson, S. E., Hoffman Bolton, J. A., Hunter, D. J., Hutchinson, A., Jacobs, K. B., Jenab, M., Khaw, K.-T., Kraft, P., Krogh, V., Kurtz, R. C., Mcwilliams, R. R., Mendelsohn, J. B., Patel, A. V., Rabe, K. G., Riboli, E., Shu, X.-O., Tjønneland, A., Tobias, G. S., Trichopoulos, D., Virtamo, J., Visvanathan, K., Watters, J., Yu, H., Zeleniuch-Jacquotte, A., Amundadottir, L. & Stolzenberg-Solomon, R. Z. 2012. Pathway analysis of genome-wide association study data highlights pancreatic development genes as susceptibility factors for pancreatic cancer. Carcinogenesis, 33, 1384–1390.

Luo, Z., Li, Y., Wang, H., Fleming, J., Li, M., Kang, Y., Zhang, R. & Li, D. 2015. Hepatocyte Nuclear Factor 1A (HNF1A) as a Possible Tumor Suppressor in Pancreatic Cancer. PLoS ONE, 10, e0121082.

Luo, Z., Li, Y., Zuo, M., Liu, C., Yan, D., Wang, H. & Li, D. 2017. Effect of NR5A2 inhibition on pancreatic cancer stem cell (CSC) properties and epithelial-mesenchymal transition (EMT) markers. Mol Carcinog, n/a-n/a.

Malakootian, M., Mirzadeh Azad, F., Naeli, P., Pakzad, M., Fouani, Y., Taheri Bajgan, E., Baharvand, H. & Mowla, S. J. 2017. Novel spliced variants of OCT4, OCT4C and OCT4C1, with distinct expression patterns and functions in pluripotent and tumor cell lines. Eur J Cell Biol, 96, 347–355.

Merlos-Suárez A., Barriga Francisco M., Jung, P., Iglesias, M., Céspedes, María V., Rossell, D., Sevillano, M., Hernando-Momblona, X., Da Silva-Diz V., Muñoz, P., Clevers, H., Sancho, E., Mangues, R. & Batlle, E. 2011. The Intestinal Stem Cell Signature Identifies Colorectal Cancer Stem Cells and Predicts Disease Relapse. Cell Stem Cell, 8, 511–524.

Miranda-Lorenzo I., Dorado, J., Lonardo, E., Alcala, S., Serrano, A. G., Clausell-Tormos J., Cioffi, M., Megias, D., Zagorac, S., Balic, A., Hidalgo, M., Erkan, M., Kleeff, J., Scarpa, A., SAINZ JR, B. & Heeschen, C. 2014. Intracellular autofluorescence: a biomarker for epithelial cancer stem cells. Nat Methods, 11, 1161–1169.

Molero, X., Vaquero, E. C., Flández, M., González, A. M., Ortiz, M. Á., Cibrián-uhalte E., Servitja, J.-M., Merlos, A., Juanpere, N., Massumi, M., Skoudy, A., Macdonald, R., Ferrer, J. & Real, F. X. 2012. Gene expression dynamics after murine pancreatitis unveils novel roles for Hnf1α in acinar cell homeostasis. Gut, 61, 1187–1196.

O’Brien C. A., Pollett, A., Gallinger, S. & Dick, J. E. 2007. A human colon cancer cell capable of initiating tumour growth in immunodeficient mice. Nature, 445, 106–110.

Ohta, T., Yamamoto, M., Numata, M., Iseki, S., Tsukioka, Y., Miyashita, T., Kayahara, M., Nagakawa, T., Miyazaki, I., Nishikawa, K. & Yoshitake, Y. 1995. Expression of basic fibroblast growth factor and its receptor in human pancreatic carcinomas. Br J Cancer, 72, 824–31.

Olive, K. P., Jacobetz, M. A., Davidson, C. J., Gopinathan, A., Mcintyre, D., Honess, D., Madhu, B., Goldgraben, M. A., Caldwell, M. E., Allard, D., Frese, K. K., Denicola, G., Feig, C., Combs, C., Winter, S. P., Ireland-Zecchini H., Reichelt, S., Howat, W. J., Chang, A., Dhara, M., Wang, L., Rückert, F., Grützmann, R., Pilarsky, C., Izeradjene, K., Hingorani, S. R., Huang, P., Davies, S. E., Plunkett, W., Egorin, M., Hruban, R. H., Whitebread, N., Mcgovern, K., Adams, J., Iacobuzio-DonahueC., Griffiths, J. & Tuveson, D. A. 2009. Inhibition of hedgehog signaling enhances delivery of chemotherapy in a mouse model of pancreatic cancer. Science, 324, 1457–1461.

Paulsen, M. T., Veloso, A., Prasad, J., Bedi, K., Ljungman, E. A., Tsan, Y.-C., Chang, C.-W., Tarrier, B., Washburn, J. G., Lyons, R., Robinson, D. R., Kumar-Sinha C., Wilson T. E. & Ljungman M. 2013. Coordinated regulation of synthesis and stability of RNA during the acute TNF-induced proinflammatory response. Proc Natl Acad Sci U S A, 110, 2240–2245.

Pierce, B. L. & Ahsan, H. 2011a. Genome-wide “pleiotropy scan” identifies HNF1A region as a novel pancreatic cancer susceptibility locus. Cancer Research, 71, 4352–4358.

Pierce, B. L. & Ahsan, H. 2011b. Genome-wide “pleiotropy scan” identifies HNF1A region as a novel pancreatic cancer susceptibility locus. Cancer Res, 71, 4352–4358.

Powell, D. & Suwanichkul, A. 1993. HNF1 activates transcription of the human gene for insulin-like growth factor binding protein-1. DNA Cell Biol, 12, 283–9.

Proctor, E., Waghray, M., Lee, C. J., Heidt, D. G., Yalamanchili, M., Li, C., Bednar, F. & Simeone, D. M. 2013. Bmi1 Enhances Tumorigenicity and Cancer Stem Cell Function in Pancreatic Adenocarcinoma. PLoS ONE, 8, e55820.

Provenzano, Paolo P., Cuevas, C., Chang, Amy E., Goel, Vikas K., Von Hoff, Daniel D. & Hingorani, Sunil R. 2012. Enzymatic Targeting of the Stroma Ablates Physical Barriers to Treatment of Pancreatic Ductal Adenocarcinoma. Cancer Cell, 21, 418–429.

Rahib, L., Smith, B. D., Aizenberg, R., Rosenzweig, A. B., Fleshman, J. M. & Matrisian, L. M. 2014. Projecting Cancer Incidence and Deaths to 2030: The Unexpected Burden of Thyroid, Liver, and Pancreas Cancers in the United States. Cancer Res, 74, 2913–2921.

Roy, N., Takeuchi, K. K., Ruggeri, J. M., Bailey, P., Chang, D., Li, J., Leonhardt, L., Puri, S., Hoffman, M. T., Gao, S., Halbrook, C. J., Song, Y., Ljungman, M., Malik, S., Wright, C. V. E., Dawson, D. W., Biankin, A. V., Hebrok, M. & Crawford, H. C. 2016. PDX1 dynamically regulates pancreatic ductal adenocarcinoma initiation and maintenance. Genes Dev, 30, 2669–2683.

Servitja, J.-M., Pignatelli, M., Maestro, M. á., Cardalda, C., Boj, S. F., Lozano, J., Blanco, E., Lafuente, A., Mccarthy, M. I., Sumoy, L., GuigÓ, R. & Ferrer, J. 2009. Hnf1α (MODY3) Controls Tissue-Specific Transcriptional Programs and Exerts Opposed Effects on Cell Growth in Pancreatic Islets and Liver. Mol. Cell. Biol., 29, 2945–2959.

Shah, R. N., Ibbitt, J. C., Alitalo, K. & Hurst, H. C. 2002. FGFR4 overexpression in pancreatic cancer is mediated by an intronic enhancer activated by HNF1alpha. Oncogene, 21, 8251–61.

Singh, S. K., Clarke, I. D., Terasaki, M., Bonn, V. E., Hawkins, C., Squire, J. & Dirks, P. B. 2003. Identification of a cancer stem cell in human brain tumors. Cancer Res., 63, 5821–5828.

Thanabalasingham, G., Shah, N., Vaxillaire, M., Hansen, T., Tuomi, T., Gašperíková, D., Szopa, M., Tjora, E., James, T. J., Kokko, P., Loiseleur, F., Andersson, E., Gaget, S., Isomaa, B., Nowak, N., Raeder, H., Stanik, J., Njolstad, P. R., Malecki, M. T., Klimes, I., Groop, L., Pedersen, O., Froguel, P., Mccarthy, M. I., Gloyn, A. L. & Owen, K. R. 2011. A large multi-centre European study validates high-sensitivity C-reactive protein (hsCRP) as a clinical biomarker for the diagnosis of diabetes subtypes. Diabetologia, 54, 2801.

Toniatti, C., Demartis, A., Monaci, P., Nicosia, A. & Ciliberto, G. 1990. Synergistic trans-activation of the human C-reactive protein promoter by transcription factor HNF-1 binding at two distinct sites. EMBO J, 9, 4467–4475.

Waghray, M., Yalamanchili, M., Dziubinski, M., Zeinali, M., Erkkinen, M., Yang, H., Schradle, K. A., Urs, S., Pasca Di Magliano, M., Welling, T. H., Palmbos, P. L., Abel, E. V., Sahai, V., Nagrath, S., Wang, L. & Simeone, D. M. 2016. GM-CSF Mediates Mesenchymal-Epithelial Crosstalk in Pancreatic Cancer. Cancer Discov.

Wang, Z., Li, Y., Kong, D., Banerjee, S., Ahmad, A., Azmi, A. S., Ali, S., Abbruzzese, J. L., Gallick, G. E. & Sarkar, F. H. 2009. Acquisition of epithelial-mesenchymal transition phenotype of gemcitabine-resistant pancreatic cancer cells is linked with activation of the notch signaling pathway. Cancer Res., 69, 2400–2407.

Wei, P., Tang, H. & Li, D. 2012. Insights into pancreatic cancer etiology from pathway analysis of genome-wide association study data. PLoS One, 7, e46887.

Yachida, S., White, C. M., Naito, Y., Zhong, Y., Brosnan, J. A., Macgregor-Das, A. M., Morgan, R. A., Saunders, T., Laheru, D. A., Herman, J. M., Hruban, R. H., Klein, A. P., Jones, S., Velculescu, V., Wolfgang, C. L. & Iacobuzio-Donahue, C. A. 2012. Clinical Significance of the Genetic Landscape of Pancreatic Cancer and Implications for Identification of Potential Long Term Survivors. Clin Cancer Res, 18, 6339–6347.

Zhu, R., Wong, K.-F., Lee, N. P. Y., Lee, K.-F. & Luk, J. M. C. 2010. HNF1α and CDX2 transcriptional factors bind to cadherin-17 (CDH17) gene promoter and modulate its expression in hepatocellular carcinoma. J Cell Biochem, 111, 618–626.

